# CRISPRa screen identifies a role for c-KIT signaling in tamoxifen resistance, potentially through upregulation of ABC transporters

**DOI:** 10.1101/2022.08.22.504845

**Authors:** Brooke A. Marks, Lauren A. Choate, Kelly Sams, Lina Zhu, Gavisha Waidyaratne, Charles G. Danko, Scott A. Coonrod

**Affiliations:** Department of Biological and Biomedical Science, College of Veterinary Medicine, Cornell University; Baker Institute for Animal Health, College of Veterinary Medicine, Cornell University; National Clinical Research Centre for Geriatric Disorders, Department of Geriatrics, Xiangya Hospital, Central South University, Changsha, Hunan, China

**Author notes:** **Author Contributions:** Manuscript preparation: Marks, B.A.; Edits: Marks, B.A., Sams K., and Coonrod, S.A.; Conception and Design: Marks, B.A., Waidyaratne G, Danko, C.G., and Coonrod, S.A. Data Interpretation: Marks, B.A., Choate L.A., Sams K., Zhu, L., Danko, C.G., and Coonrod, S.A.; Experiments: Marks, B.A.

## Abstract

Resistance to endocrine therapy is a common problem in patients with estrogen receptor alpha (ER*α*) positive breast cancer. In this study, we took a non-biased genome-wide approach to identify novel mechanisms of endocrine resistance using a clustered regularly interspaced short palindromic repeats (CRISPR) activating (CRISPRa) screen. Results from the screen identified 109 candidate resistance-associated genes, with several of these genes, such as EGFR and SRC, having been previously associated with endocrine resistance. One candidate gene that has not been previously associated with endocrine resistance is the tyrosine kinase receptor, c-KIT. We further tested for associations between c-KIT and endocrine resistance and found that c-KIT overexpressing cells proliferate more rapidly in the presence of tamoxifen compared to control cell lines. To gain deeper insight into the potential role of c-KIT signaling in tamoxifen resistance, we next performed precision nuclear run-on and sequencing (PRO-seq) analysis of c-KIT overexpressing cells to identify downstream factors that may mediate the c-KIT response. Kyoto Encyclopedia of Genes and Genomes (KEGG) analysis of the overexpressed genes found that the only class of factors that was significantly induced by c-KIT was the ATP-binding cassette (ABC) transporters; specifically, ABCA1, ABCA4, and ABCG1. Interestingly, overexpression of two of these ABC transporters, ABCA1 and ABCG1, significantly correlated with worse prognosis in ER*α*+ breast cancer patients following endocrine therapy. We then tested for potential therapeutic effects of c-KIT inhibition on endocrine resistance and found that the c-KIT inhibitor Gleevec appears to synergize with tamoxifen to suppress MCF-7-S cell growth. Together, our findings support the hypothesis that c-KIT signaling promotes endocrine resistance via the induction of ABC transporter activity. Additionally, our studies suggest that inhibition of c-KIT signaling may represent a novel strategy for preventing or overcoming endocrine resistance in ER*α*+ patients.

## Background

Endocrine therapies such as tamoxifen (TAM) are currently used to treat estrogen receptor alpha positive (ER*α*+) breast cancer patients. ER*α*+ breast cancers (BC) utilize estrogen signaling to proliferate and survive, and endocrine therapies suppress tumor growth and spread by blocking estrogen signaling. The use of these therapies, as well as mammography screening, has led to a decline in the number of breast cancer deaths over the years (Berry et al., 2005). However, despite these advances, between 20-30% of patients have *de novo* or acquired resistance to these therapies, leading to tumor progression, metastasis, and increased mortality rates (Ali et al., 2016). Additionally, up to 50% of patients with metastatic disease will not respond to endocrine therapy as the first line of treatment (Anderson, Bulun, Smith, & Dowsett, 2007).

Therefore, there is a pressing need to develop additional treatments that can be used, either alone or in combination with current therapies, to treat endocrine resistant patients. Outcomes from ongoing studies have facilitated the development of FDA approved CDK 4/6 inhibitors (abemaciclib, palbociclib, and ribociclib), used to treat metastatic BC patients, and the mTOR inhibitor, everolimus, which is approved to treat postmenopausal ER*α*+ BC patients. Additionally, promising therapeutics have also been developed that target other pathways implicated in endocrine resistance including, EGFR/HER2 (Jiang et al., 2014; Massarweh et al., 2008; TW et al., 2009; Zhang et al., 2015), IGF-1R (Fox et al., 2011; TW et al., 2009), FGFR (Drago et al., 2019; Mouron et al., 2021), and RET (Morandi et al., 2013) signaling. Despite these recent advances, it seems likely that additional endocrine resistance mechanisms remain to be defined.

Therefore, to gain further insight into the mechanisms of resistance, we performed an unbiased, genome wide CRISPR activating (CRISPRa) screen to identify genes that, when overexpressed, enhanced the survival of tamoxifen-sensitive MCF-7 (MCF-7-S) cells in the presence of TAM. To date, most genome-wide screens have used knockdown approaches to identify endocrine resistance genes in BC cell lines (Ahn et al., 2010; Fox et al., 2011; Mendes-Pereira et al., n.d.). While knockdown screens are extremely useful, this method has the specific limitations of only identifying factors whose *absence* promotes endocrine resistance and/or identifying the active resistance pathway(s) in the cell line being investigated. The CRISPRa screen avoids these limitations by inducing target gene transcription of most known protein coding genes, independent of parent cell line transcription levels, to investigate the involvement of specifically upregulated genes in TAM resistance. Moreover, our approach may have an additional benefit over these knockdown approaches because the CRISPRa screen should identify factors that promote resistance when overexpressed, thus more closely resembling the oncogenic environment that occurs during the acquisition of endocrine resistance, as observed when signaling pathways, such as EGFR/HER2 are upregulated.

Results from our screen identified a number of genes that have been previously implicated in endocrine resistance, such as EGFR (Jiang et al., 2014; Massarweh et al., 2008; TW et al., 2009; Zhang et al., 2015) and SRC (Guest et al., 2016; Hiscox, Barrett-Lee, Borley, & Nicholson, 2010). However, we also found that other genes, including the tyrosine kinase receptor c-KIT (also known as stem cell factor receptor, KIT, or CD117) appeared to promote resistance. c-KIT signaling is known to be important for hematopoietic stem cell maintenance (Kimura et al., 2011) and also plays a role in certain cancers such as gastrointestinal stromal tumors (GIST) (M & J, 2001), acute myeloid leukemia (Schwartz et al., 2009), small cell lung cancer (Naeem, Dahiya, Clark, Creech, & Alkan, 2002), glial cancer (Cetin, Dienel, & Gokden, 2005), and melanoma (Pham, Guhan, & Tsao, 2020). c-KIT has also been linked, albeit with conflicting results, to breast cancer (Noronha et al., 2011; Simon et al., 2004; Tsuda et al., 2005). To date, however, c-KIT has yet to be implicated in endocrine resistance. Our analysis of c-KIT in this context demonstrated that its overexpression does appear to promote endocrine resistance in cell models, potentially via the upregulation of ABC transporter proteins. Lastly, we found that the c-KIT inhibitor, Gleevec, appears to synergize with TAM in blocking MCF-7-S cell growth, suggesting that c-KIT inhibition may represent a new target for endocrine therapy.

## Results

### Identification of TAM resistant candidate genes using a CRISPR activating (CRISPRa) screen

To identify novel genes important in the development of TAM resistance, we generated an MCF-7-S cell line that contained two stably integrated components: (a) a tetracycline (TET)-inducible dCAS9-VP64 construct (Figure 1B-D) and (b) a constitutively expressed CRISPRa sgRNA library that targets most known protein-coding gene promoters (Gilbert et al., 2014) (Figure 1A and E). The library (Gilbert et al., 2014; Horlbeck et al., 2016) contains a total of 104,540 sgRNAs (including non-targeting sgRNAs) with each gene promoter being targeted by 5 separate sgRNAs from the library. To prepare for the screen, MCF-7-S cells containing the integrated inducible dCAS9-VP64 construct, were transduced with lentiviruses containing the sgRNA library at a low infection rate to obtain approximately one sgRNA per cell (Kampmann, Bassik, & Weissman, 2014). Following sgRNA integration into the genome, cells were selected using puromycin. Cells were then treated with vehicle or doxycycline (DOX, a TET derivative) for 24hrs to induce dCAS9 and subsequent target gene transcription, followed by twenty days of treatment with or without TAM (Figure 1E). Cells from the four different treatment groups (-DOX/-TAM, -DOX/+TAM, +DOX/-TAM, and +DOX/+TAM) were harvested and prepared for Illumina sequencing (Figure 1E). The integrated sgRNA DNA in each cell was then sequenced and then computationally linked with its specific targeted gene. Note that sgRNAs are identified by their target genes’ names in this report. The sgRNAs that had differences in abundance between the four treatment groups were then identified using DESeq2 (Figure 1E, Figure 2A-B, and Figure S2A-B) using an FDR of < 0.05 and a fold enrichment of ≥ 2.5.

**Figure 1.**
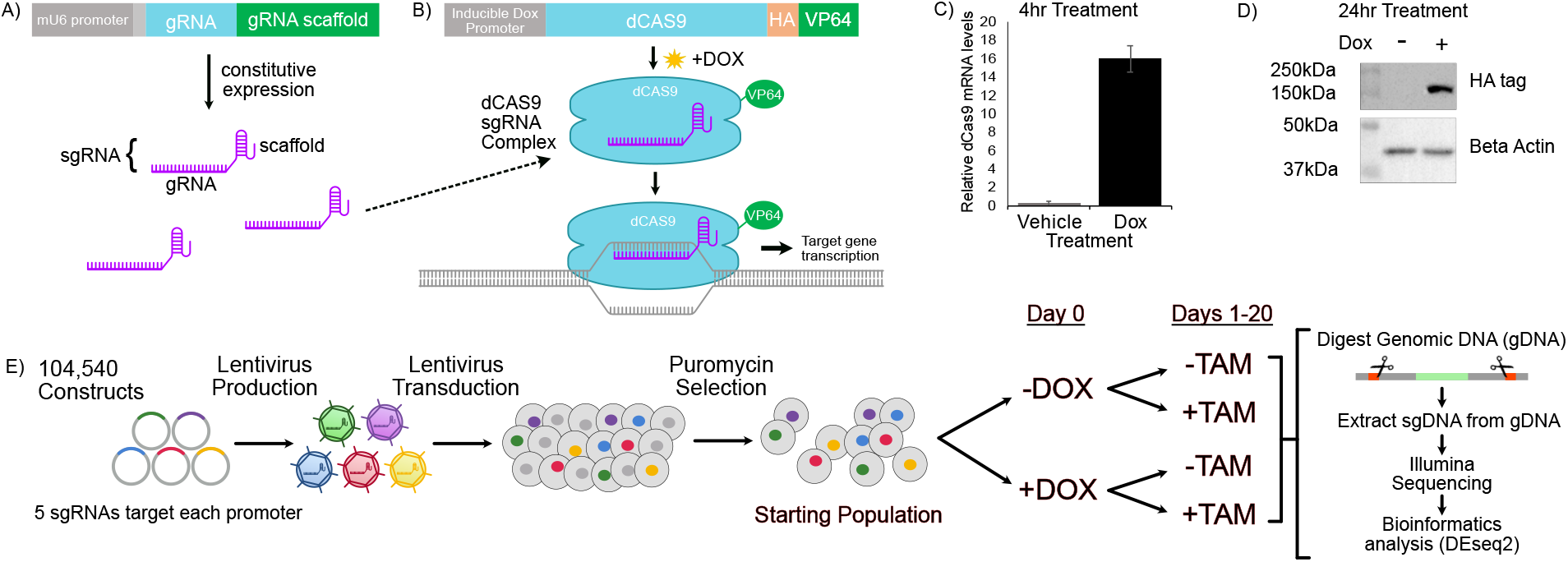
Schematic representation of methods to identify tamoxifen resistant candidate genes using a CRISPR activating (CRISPRa) screen. Schematic of the (A) sgRNA and (B) doxycycline-inducible dCAS9 activating system used in MCF-7-S cells. C) Relative dCAS9 mRNA transcription within the MCF-7-S dCAS9-VP64 stable cell line was increased 16-fold after 4hrs of DOX treatment when compared to vehicle treatment. D) The HA-dCAS9-VP64 fusion protein was strongly expressed after 24hr of DOX treatment compared to vehicle treatment in MCF-7-S dCAS9-VP64 cells. E) Schematic representation of CRISPRa screen methodology.

**Figure 2.**
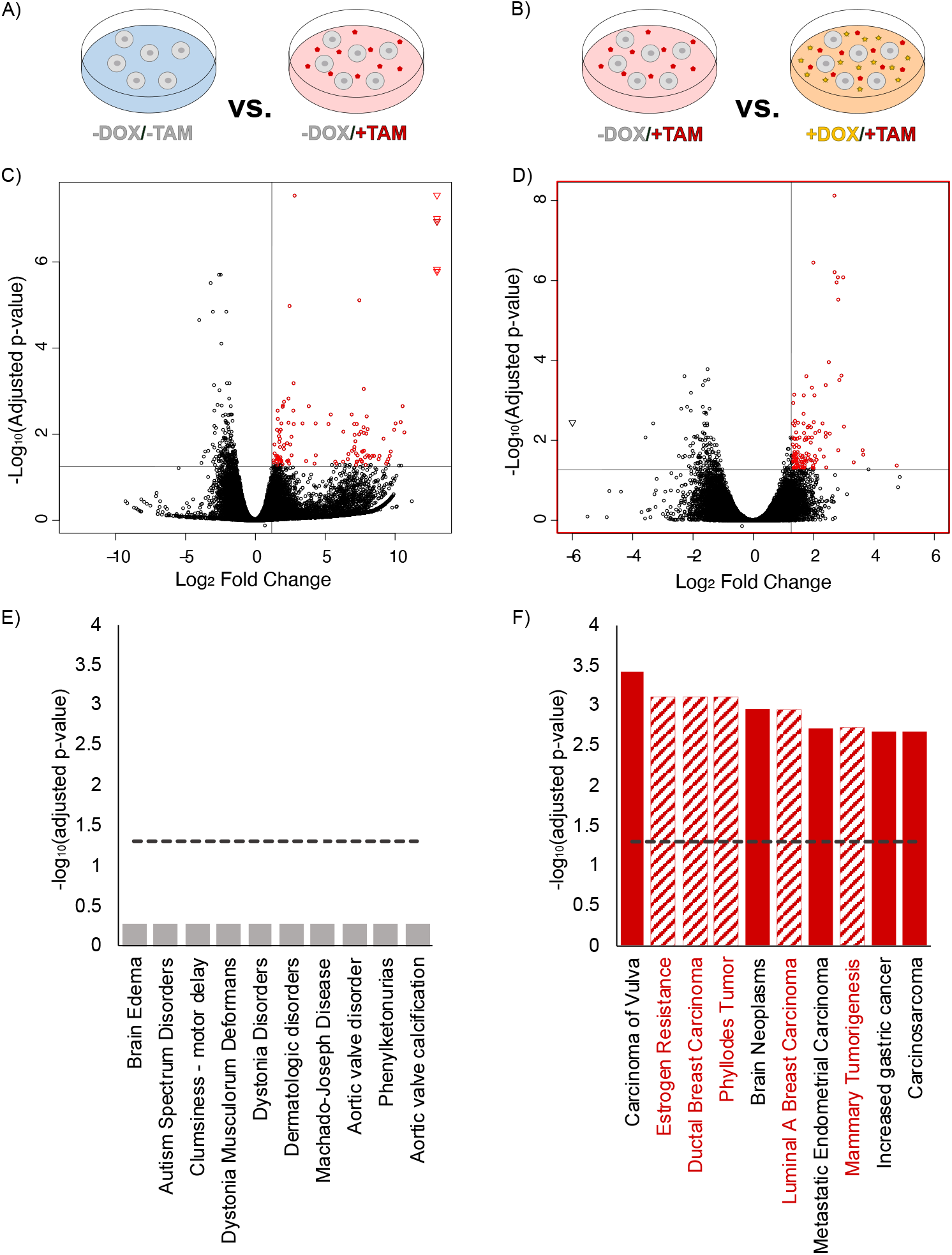
Analysis of CRISPRa screen identifies candidate genes associated with BC and resistance in MCF-7-S cells. A-B) Schematic representation of treatment conditions corresponding to the volcano plots and bar charts shown below them. C) Volcano plot in which the upper right quadrant identifies sgRNAs (genes) that were significantly over-represented when comparing cells in the absence of any CRISPR induction (-DOX) and in the presence of TAM (-DOX/+TAM) to those without CRISPR induction or TAM treatment (-DOX/-TAM), (p < 0.05). D) Volcano plot in which the upper right quadrant identifies sgRNAs that were significantly over-represented in cells with CRISPR induction and TAM treatment (+DOX/+TAM) compared to cells with no induction and TAM treatment (-DOX/+TAM), (p < 0.05). Circles represent individual sgRNAs, and outliers are represented as triangles. E-F) DISGENET disease enrichment database (within the ENRICHR database) was used to identify the top 10 human diseases associated with the treatment specific subsets of enriched sgRNAs (genes) identified in the volcano plots above, showing that a majority of the sgRNAs in our experimental group (F) that appear to increase cell survival in the presence of tamoxifen have been previously implicated in breast cancer and resistance (observed as striped bars). Dotted line represents the adjusted p-value < 0.05.

Comparison of the treatment groups found that 115 sgRNAs (genes) were observed to have differences in abundance between the -DOX/-TAM group and the -DOX/+TAM group (Figure 2C) and 75 sgRNAs (genes) had abundance differences between the -DOX/-TAM group and the +DOX/-TAM group (Figure S2C). One hundred and ninety-seven sgRNAs (genes) showed differences in abundance between the +DOX/-TAM and +DOX/+TAM treatment group (Figure S2D). Lastly, we identified 109 sgRNAs (genes) that had differences in abundance between the -DOX/+TAM and +DOX/+TAM treatment groups (Figure 2D). We considered – DOX/+TAM and +DOX/+TAM to be our experimental comparison because we predicted that, following DOX treatment, cells expressing genes that promote TAM resistance would continue to proliferate in the presence of TAM when compared to the cells that were treated with TAM but were not treated with DOX (Figure 2B).

To begin analyzing the sets of genes that were identified through differential expression between the treatment groups, we ran these gene lists through the DisGeNET database (Piñero et al., 2017) within the ENRICHR database (EY et al., 2013; MV et al., 2016; Xie et al., 2021). The DisGeNET database contains collections of genes associated with specific human diseases and output from this analysis provides information on the disease, the corresponding genes associated with the disease, and the adjusted p-value. Results from this analysis showed that the – DOX/-TAM vs. -DOX/+TAM and the +DOX/-TAM vs. +DOX/+TAM comparisons did not contain any significantly enriched disease datasets according to the adjusted p-value (p < 0.05). The -DOX/-TAM vs. +DOX/-TAM comparison contained four significantly enriched disease datasets, however, none of these datasets were associated with breast cancer or hormone resistance. Interestingly, we found that the experimental comparison (-DOX/+TAM vs. +DOX/+TAM) dataset showed a significantly higher association with specific diseases. Importantly, of the top ten enriched disease gene sets identified in this group (Figure 2F, Table S9), five of these sets were associated with breast cancer, with one of these sets being specifically associated with estrogen resistance. This observation suggests that our CRISPRa screen was, in fact, enriching for endocrine resistance genes, and our results also suggest that genes identified in the experimental comparison represent strong candidates for further study.

### Candidate gene signaling pathways

Given the previous observation, we decided to further investigate the involved pathways corresponding to the significant sgRNAs (genes) in the experimental dataset (-DOX/+TAM vs. +DOX/+TAM, Figure 3A, Table S10) using the ‘enrichment map’ function in the R package ‘HTSanalyzeR2’ (L.Zhu, F. Gao, X. Mei, n.d.). This enrichment map uses a gene over-representation analysis to compare gene ontologies (GO) and signaling pathways from known databases to a specific input gene list (experimental dataset; -DOX/+TAM vs. +DOX/+TAM) and visualizes GO and signaling pathway categories that are over-represented in the input list. Results show that two of the gene sets identified in this analysis were “breast cancer” and “endocrine resistance” (Figure 3A), thus supporting the outcomes from the DisGeNET analysis. Other signaling pathways involved in cancer identified in this analysis included EGFR, HER2, MAPK, PI3K-AKT, and JAK-STAT signaling. Importantly, EGFR signaling through the MAPK pathway has long been associated with both experimental and clinical endocrine therapy resistance (Jiang et al., 2014; Massarweh et al., 2008; TW et al., 2009; Zhang et al., 2015).

**Figure 3.**
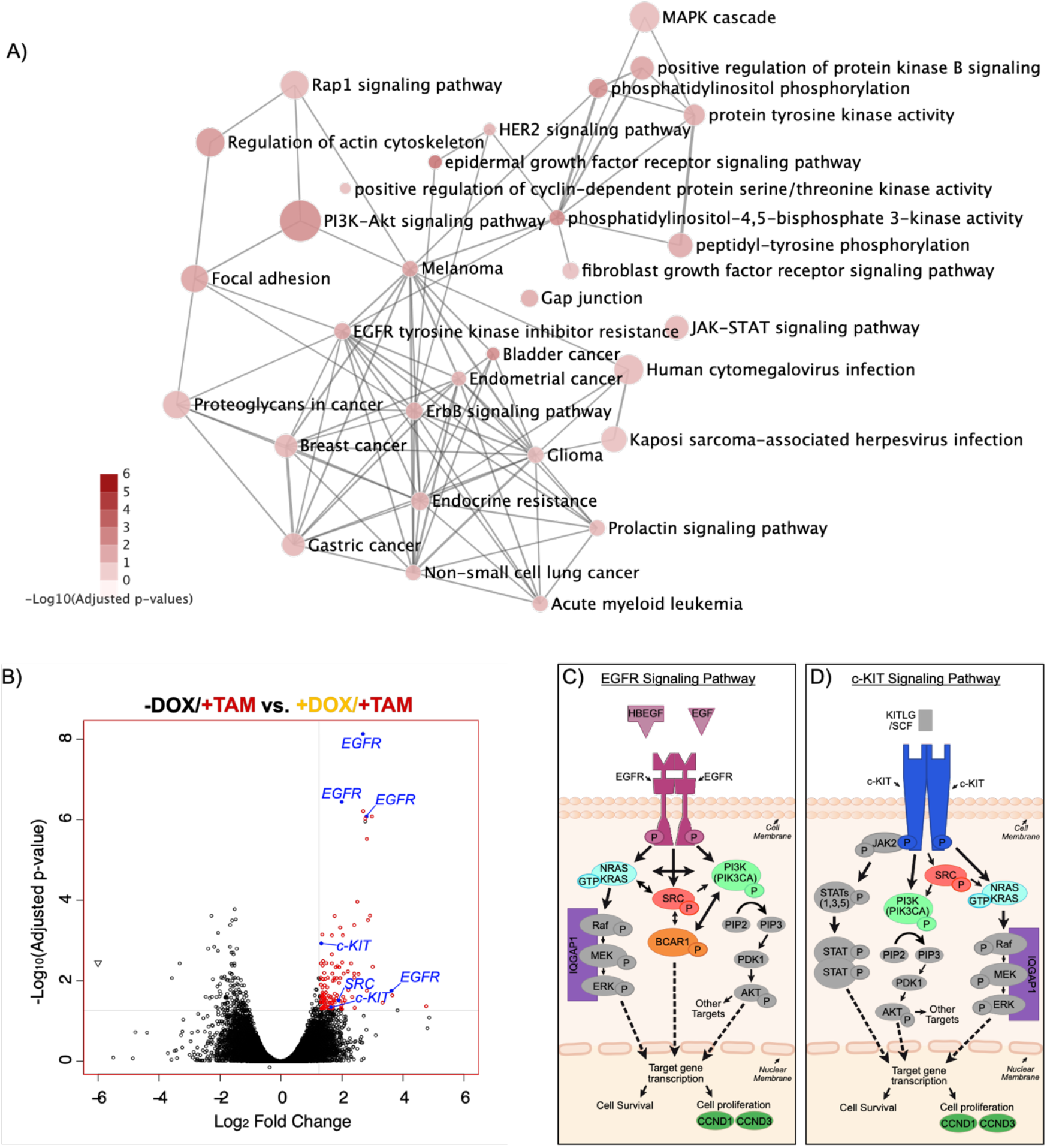
TAM resistant candidate genes (sgRNAs) are enriched in multiple cancers and cancer signaling pathways. A) The enrichment map shows signaling pathways and gene sets enriched in TAM resistant candidate genes identified by ‘HTSanalyzeR2’. Circle size increases with the number of genes associated with a specific pathway, and the shade of red increases with statistical significance. B) Specific resistant candidate genes (sgRNAs) observed in the volcano plot of -DOX+TAM vs. +DOX+TAM comparison. Red dots are significantly enriched (p < 0.05 and fold enrichment > 2.5) and blue dots are known resistance genes (EGFR, SRC) and the candidate TAM resistant gene (c-KIT). C) EGFR and D) c-KIT downstream signaling components. Genes (sgRNAs) represented at least once in the experimental dataset (Table S5) are shown in color and unidentified components are represented in grey.

A more detailed view of the experimental group (-DOX/+TAM vs. +DOX/+TAM) volcano plot (Figure 3B) shows that, four of five EGFR sgRNAs used in the screen and represented as the gene name EGFR, were significantly enriched. Given how closely the EGFR tyrosine kinase receptor has been linked to endocrine resistance (Jiang et al., 2014; Massarweh et al., 2008; TW et al., 2009; Zhang et al., 2015), this result further strengthens the hypothesis that enriched sgRNAs (candidate genes) identified in our screen represent strong candidates for promoting endocrine resistance. Interestingly, we also found that two of the five c-KIT sgRNAs (represented as the gene name c-KIT) were significantly enriched in our screen(Figure 3B). As opposed to EGFR, the c-KIT tyrosine kinase receptor has not been previously implicated in endocrine resistance. Given the novelty of this observation, we then decided to further test for potential links between c-KIT and endocrine resistance. We also included EGFR in our subsequent validation studies as a positive control.

We next evaluated the list of genes that were differentially expressed in our experimental group comparison (-DOX/+TAM vs. +DOX/+TAM) to see if additional factors previously associated with either EGFR or c-KIT signaling were contained within the list (Figure 3C-D). Results show that several factors that are downstream of both signaling pathways were identified in our screen. These factors include, NRAS, PIK3CA, IQGAP1, SRC, CCND1, and CCND3. Some factors have also been previously associated with one pathway but not the other. For example, BCAR1 is a known downstream target of the EGFR signaling pathway but not a known downstream target of c-KIT signaling. Additionally, both HBEGF and EGF were identified in the screen and these two factors are known ligands for EGFR, however, they have not been associated with c-KIT signaling. Interestingly, SRC, which was significantly enriched in the screen and is associated with both signaling pathways (Figure 3B-D), has been associate with a range of cancers [reviewed in (Irby & Yeatman, 2000; Wheeler, Iida, & Dunn, 2009)], including BC and has also been associated with endocrine resistance (Guest et al., 2016; Hiscox et al., 2010). Given these strong associations, we also included SRC as a positive control in the validation studies below. Several other components linked to these signaling pathways were not identified in our screen (show in gray). This could be because either 1) the activity of these factors is dependent upon activation (phosphorylation etc.) as opposed to overexpression or 2) the factors identified are the rate limiting factors in the pathway, while the unidentified factors are already over-abundant in the cell.

### Validation of selected genes

We next carried out cell proliferation assays to more directly test whether MCF-7-S cell lines that specifically overexpress (OE) either EGFR (MCF-7-S^EGFR-OE^), SRC (MCF-7-S^SRC-OE^), or c-KIT (MCF-7-S^c-KIT-OE^) are endocrine resistant. These three cell lines were generated using the same inducible dCAS9-VP64 construct and the sgRNAs that were significantly upregulated in the screen, as described in Figure 1. After the development of these cell lines, we first performed quantitative PCR (qPCR) analysis of the target genes to quantify mRNA expression levels after 0hr-48hrs of DOX treatment to induce gene expression. Results show that the expression of EGFR (Figure 4A), SRC (Figure 4B), and c-KIT (Figure 4C) were induced in a time-dependent manner in the respective MCF-7-S^EGFR-OE^, MCF-7-S^SRC-OE^, and MCF-7-S^c-KIT-OE^ cell lines.

**Figure 4:**
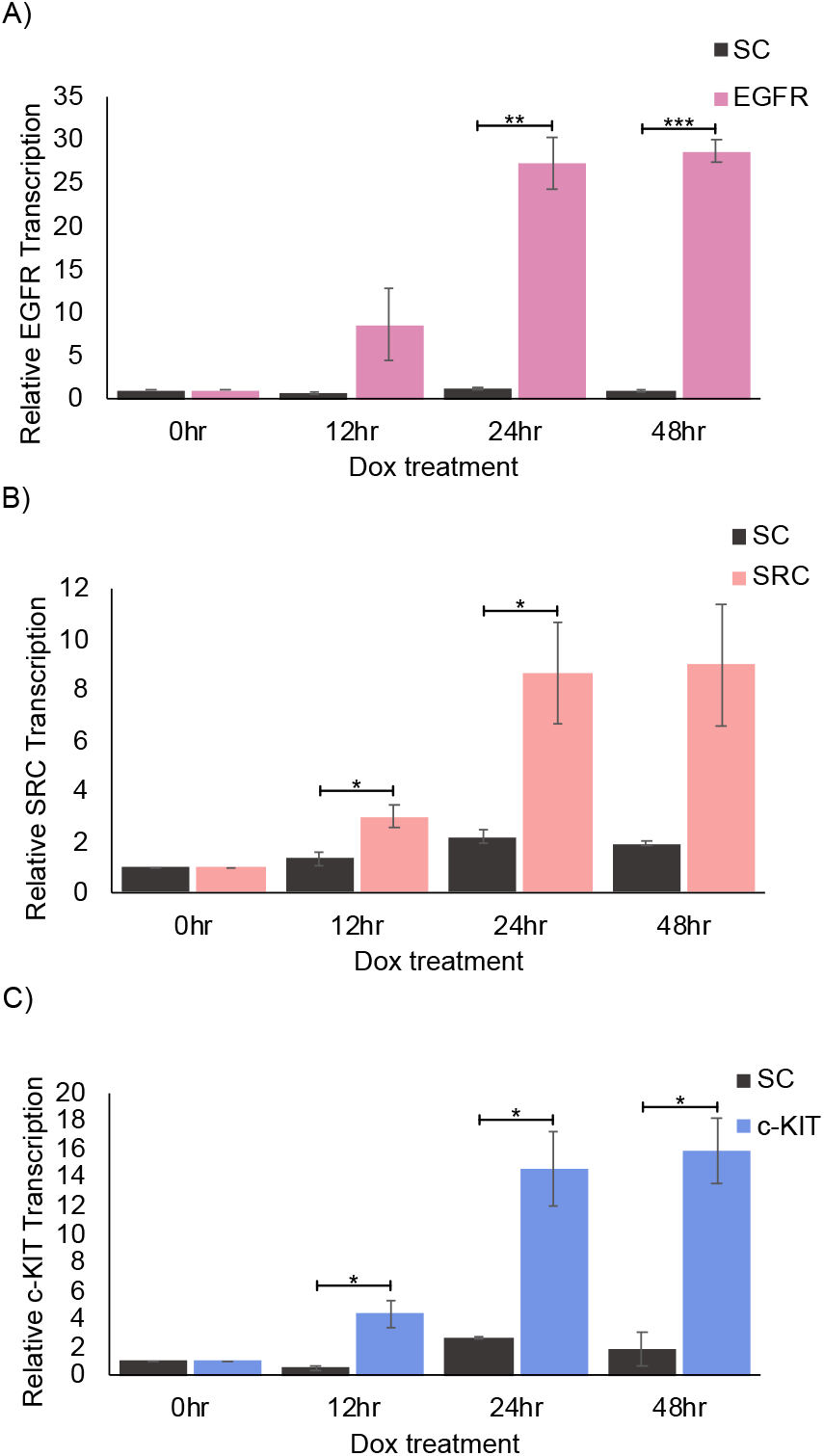
Target gene upregulation in MCF-7-S cells. Relative mRNA transcription levels of dox inducible A) MCF-7-S^EGFR^, B) MCF-7-S^SRC^, and C) MCF-7-S^c-KIT^ overexpressing lines after DOX treatment. * p < 0.05. ** p < 0.01. *** p < 0.005.

To evaluate cell proliferation and the importance of these specific genes in resistance, the cell lines were treated with or without DOX for 24hrs and then treated with or without DOX and with TAM for a minimum of 5 days or until cells were close to 100% confluency (Figure 5A). We used this treatment regime to recapitulate the screening procedure as closely as possible. Analysis of cell proliferation was performed using the equation shown in 5B. This type of analysis allowed for normalization of the experiment based on differences in growth. Results show that EGFR, SRC, and c-KIT overexpressing cells proliferated more rapidly in the presence of TAM compared to both the scrambled (SC) and random gene (A2M) controls (Figure 5C). Taken together, these proliferation studies provide further support for the hypothesis that EGFR, SRC, and c-KIT play a role in endocrine resistance.

**Figure 5.**
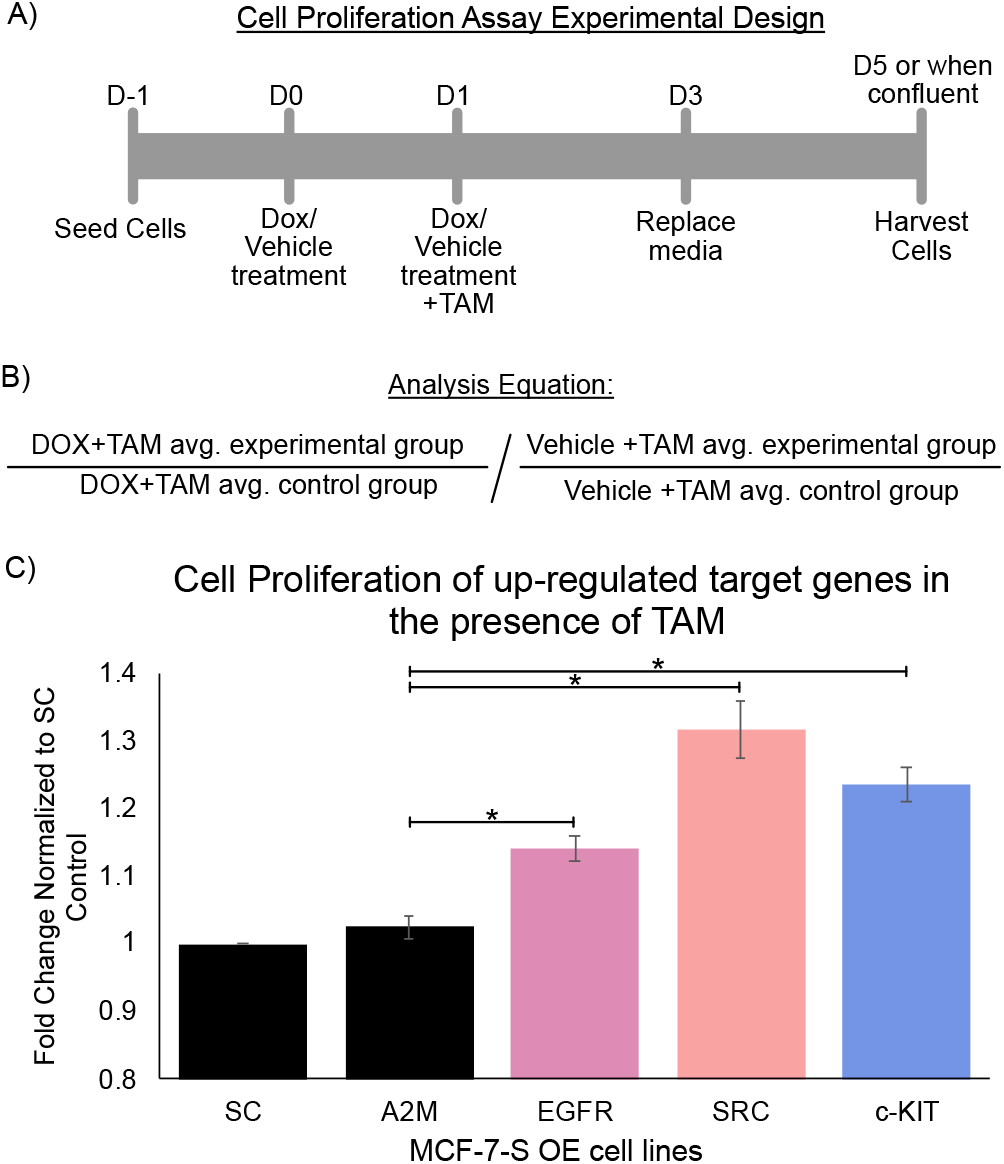
Upregulation of EGFR, SRC, and c-KIT individually promote TAM resistance. A) Experimental timeline of cell line treatments. B) Equation analysis used to determine cell proliferation in the presence of TAM. C) Cell proliferation rates of MCF-7-S^EGFR^, B) MCF-7-S^SRC^, and C) MCF-7-S^c-KIT^ overexpressing cells were increased compared to MCF-7-S^SC^and MCF-7-S^A2M^ (gene control) czlls when in the presence of TAM.

### Identification of a potential resistance mechanism in c-KIT induced cells using PRO-seq

In order to begin investigating the potential mechanisms by which c-KIT promotes endocrine resistance, we next utilized precision nuclear run-on and sequencing (PRO-seq) (Mahat et al., 2016) to identify downstream factors that may be regulated by c-KIT signaling (Figure 6A). We first tested the specificity of the c-KIT sgRNA-dCAS9-VP64 complex in activating c-KIT transcription by treating MCF-7-S^c-KIT-OE^ cells with either vehicle or DOX for 24hrs (Figure 6B). Results show that c-KIT expression was potently upregulated in the DOX treated cells compared to vehicle treated cells. These results demonstrate that our CRISPR system can upregulate the expression of c-KIT with a high degree of specificity. Two other genes, CENPT and SLC25A36, were also significantly upregulated after DOX treatment. Neither of these genes has been specifically associated with c-KIT signaling, and further investigation would be needed to determine if it is directly due to c-KIT upregulation. Next, to begin investigating the mechanism by which c-KIT promotes TAM resistance, we pre-treated cells with vehicle or DOX for 24hrs and then treated the cells with TAM for 5 days, leading to two treatment groups (vehicle/+TAM vs. DOX/+TAM) (Figure 6A). We then performed PRO-seq analysis of the samples to measure transcriptional activity and identified differentially expressed genes using DESeq2 (Figure 6C). We identified 70 differentially expressed genes when comparing vehicle/+TAM with DOX/+TAM using a p-value of p < 0.075, (Table S11) and these genes were then analyzed using KEGG 2021 (Kanehisa, 2019; Kanehisa, Furumichi, Sato, Ishiguro-Watanabe, & Tanabe, 2021; Kanehisa & Goto, 2000) in the ENRICHR database (EY et al., 2013; MV et al., 2016; Xie et al., 2021). Results showed that the ATP-binding cassette (ABC) transporter genes (ABCA1, ABCA4, ABCG1) were the only functional group to be significantly enriched in the KEGG 2021 database using the p < 0.075 cutoff value (Figure 6D).

**Figure 6.**
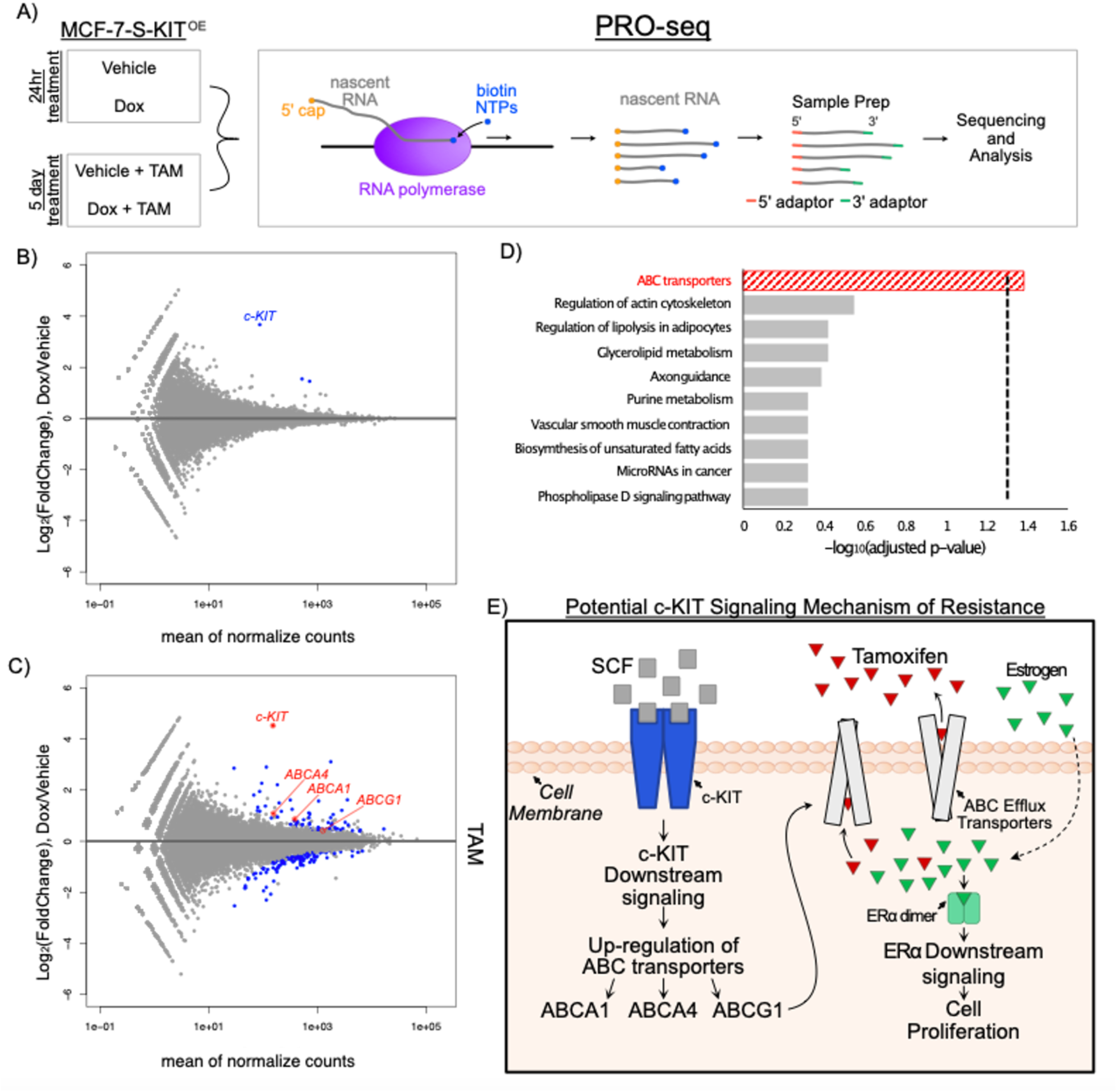
Overexpression of c-KIT in inducible MCF-7-S^c-KIT^ cells upregulates ABC transporters in the presence of TAM. A) Experimental setup for PRO-seq. PRO-seq libraries were prepared after 24hrs of vehicle or DOX treatment and after 5 days of vehicle + TAM and DOX + TAM treatment in MCF-7-S^c-KIT^ cells. B) MA plot results showing significantly upregulated genes in blue after 24hrs of DOX treatment when compared to vehicle treatment. C) MA plot results showing changed gene transcription (p < 0.075) in blue after 5 days of DOX + TAM treatment compared to vehicle + TAM treatment. Genes upregulated after DOX + TAM treatment are observed in the top half of the MA plot. Labeled genes in the plots are c-KIT and genes in the ABC transporter family. D) KEGG 2021 Human database results, from ENRICHR database, using enriched genes from figure 6C (Table S12) indicates the top 10 gene function sets. Significant gene functions, using adjusted p-value of < 0.05 (dotted line), are represented by striped bars. E) Proposed working hypothesis of resistance mechanism following activation of c-KIT signaling in c-KIT positive breast cancer cells.

### High expression of ABCA1 and ABCG1 leads to a worse prognosis in ER*α*+ BC patients treated with endocrine therapy

As a test of the hypothesis that ABC transporters play a clinically relevant role in endocrine resistance in breast cancer patients, we performed a Kaplan-Meier (Lánczky & Győrffy, 2021) plot analysis on the ABCA1, ABCA4, and ABCG1 genes for relapse-free survival (RFS) in ER*α*+ patients who had received previous endocrine therapy. We found that high expression of ABCA1 and ABCG1 are both significantly correlated with worse prognosis in ER*α*+ breast cancer patients who were treated with endocrine therapy compared to patients with low expression of ABCA1 or ABCG1 (Figure 7A-B). ABCA4 expression was not correlated with a worse prognosis (data not shown).

**Figure 7.**
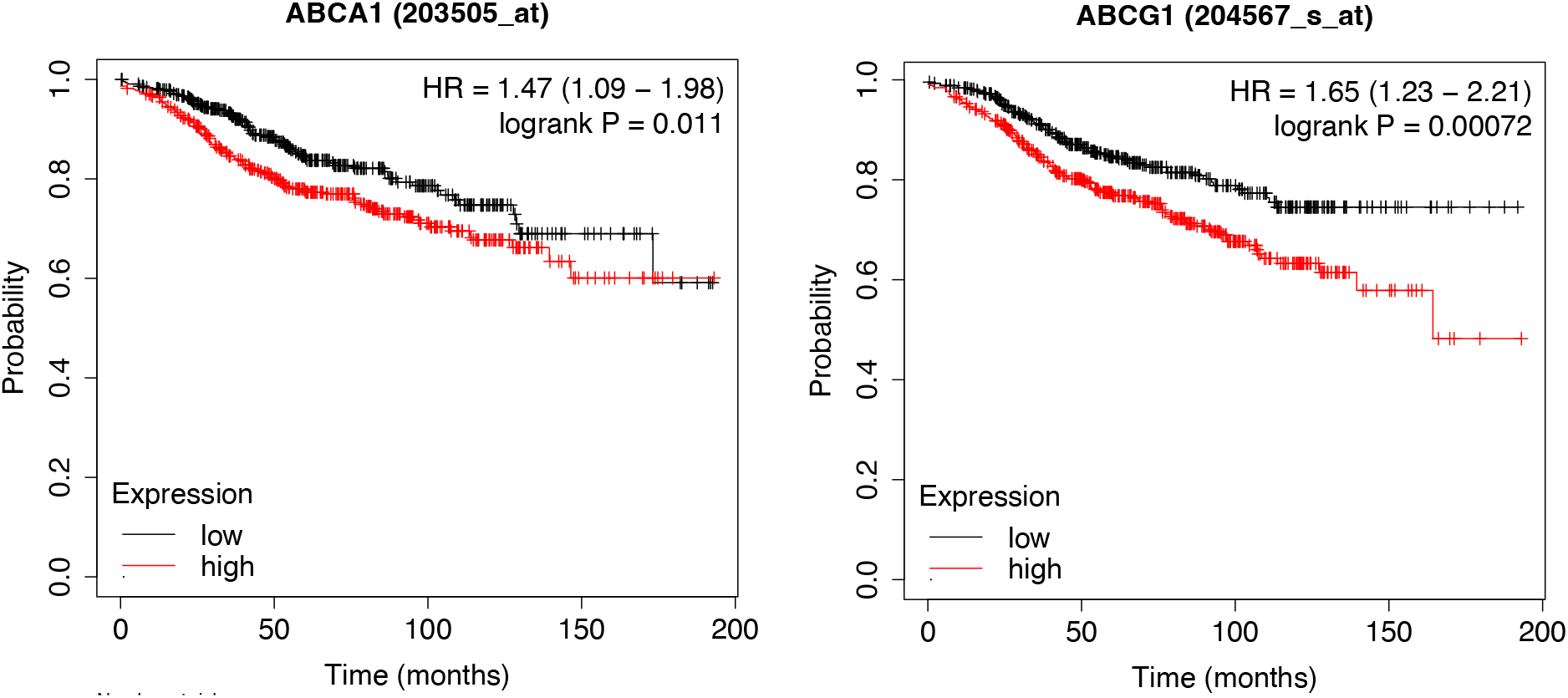
Two c-KIT downstream ABC transporters predict response free survival (RFS) in human BC patients treated with endocrine therapy. ER*α*+ BC patients with high expression of A) ABCA1 or B) ABCG1 after treatment with endocrine therapy have decreased RFS compared to patients with low expression of either ABCA1 or ABCG1.

### TAM and Gleevec synergize to suppress MCF-7-S cell growth

Given our findings linking c-KIT with endocrine resistance, we next sought to test whether pharmacological inhibition of c-KIT signaling may synergize with TAM to suppress MCF-7-S cell growth. c-KIT tyrosine kinase inhibitors such as imatinib mesylate (Gleevec) are currently being used to treat c-KIT-positive gastrointestinal stromal tumors, acute lymphoblastic, leukemia, and chronic myeloid leukemia. To determine if a TAM and Gleevec combination therapy could have an additive or synergic effect on the inhibition of MCF-7-S cells, we performed a checkerboard assay and used the SynergyFinder version 2.0 (Ianevski, Giri, & Aittokallio, 2020) to investigate cell viability and synergism after drug treatment. We found that individual treatment of MCF-7-S cells with Gleevec and TAM inhibited cell proliferation (Figure 8A). Moreover, increasing concentrations of Gleevec and TAM in combination resulted in greater inhibition of MCF-7-S cell growth compared to individual treatment (Figure 8A). Additionally, analysis using the Zero Interaction Potency (ZIP) program in SynergyFinder 2.0 to determine synergy, identified an additive and/or synergistic effect at 3.125uM, 6.25uM, and 12.5uM Gleevec in combination with 0.25uM-4uM TAM (Figure 8B). This data suggests that dual combination treatment with TAM and Gleevec could be a more beneficial treatment option for ER*α*+ breast cancer patients who also express c-KIT.

**Figure 8.**
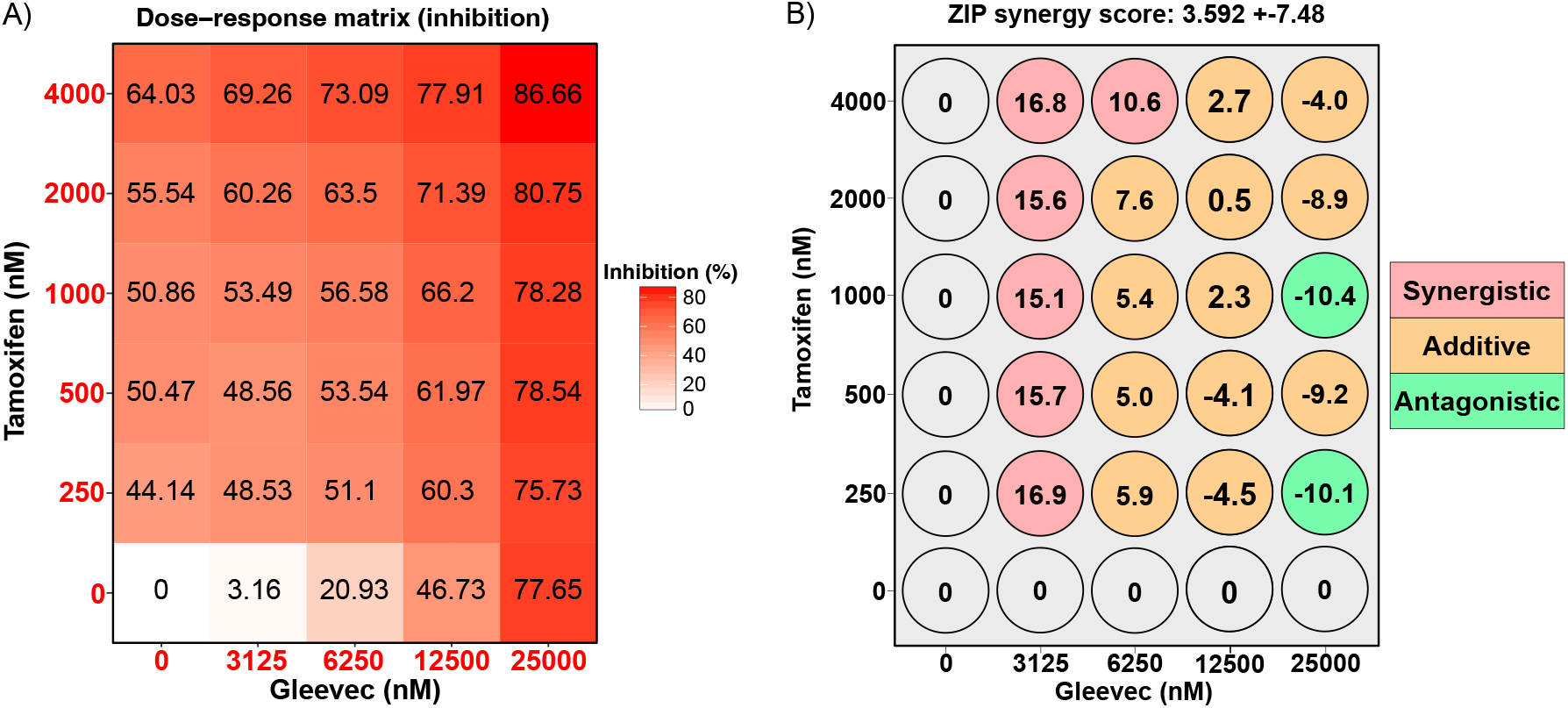
TAM and Gleevec combination treatment results in greater inhibition and drug synergism in MCF-7-S cells. A) The percentage of inhibition of MCF-7-S cells treated with increasing concentrations of TAM and Gleevec. The shade of red represents the amount of inhibition in cells. B) Observed synergism scores that correspond to the percent inhibition values (Figure 8A) of TAM and Gleevec treatment using ZIP synergy from SynergyFinder 2.0.

## Discussion

Understanding the aberrant mechanisms that drive breast cancer progression and resistance are important aspects of cancer research and this knowledge has led to the development of targeted therapies, such as tamoxifen. Unfortunately, resistance to targeted therapies is a common occurrence and understanding the molecular changes that lead to acquired and *de novo* endocrine resistance will be a critical component to developing new patient therapies. To gain insight into factors that may promote resistance, previous genome-wide screens have primarily utilized knockdown approaches. In our study, the induction of the expression of individual genes in single cells represents an alternate approach to identify resistance mechanisms and may be more representative of the gain-of-function types of mechanisms that occur in cancer patients.

Results from our CRISPRa screen identified 109 differentially expressed genes in our experimental comparison group (p < 0.05, fold enrichment ≥ 2.5). When inputting these genes through the DisGeNET database in the ENRICHR system, five out of the top ten significantly enriched disease processes were linked to breast cancer and estrogen resistance. Moreover, this dataset also identified endometrial cancer, another hormonally driven cancer, as an enriched disease process. Importantly, of the four different comparisons analyzed in our CRISPRa screen, our experimental group was the only group that contained enriched disease processes for breast cancer and estrogen resistance, suggesting that genes within this dataset are likely candidates for promoting endocrine resistance.

Further investigation into specific signaling pathways within our experimental dataset using an HTSanaylzeR2 enrichment map (L.Zhu, F. Gao, X. Mei, n.d.) found that there was an enrichment of factors that are associated with both breast cancer and endocrine resistance in this group. Moreover, this analysis also identified signaling pathways previously known to be important in cancer signaling, including the MAPK cascade, PI3K-Akt signaling, JAK-STAT signaling, ERBB2 signaling, and EGFR signaling. EGFR is associated with endocrine resistance both experimentally and clinically (Jiang et al., 2014; Massarweh et al., 2008; TW et al., 2009; Zhang et al., 2015) and the finding that four of the five EGFR sgRNAs in our library were significantly enriched in our experimental group dataset, provides further support that our screen significantly enriched for genes that play a role in endocrine resistance and also further highlights the importance of EGFR signaling in endocrine resistance. Additionally, we found that SRC was significantly enriched in our screen and that SRC appeared multiple times in the signaling pathways identified in the HTSanaylzeR2 enrichment map. This result is not surprising given that SRC is a known downstream target in multiple tyrosine kinase signaling mechanisms (including EGFR) and has been implicated in multiple cancers (Wheeler et al., 2009).

SRC is also a downstream target of the tyrosine kinase receptor c-KIT and we identified two c-KIT sgRNAs in our screen. The c-KIT signaling pathway has been implicated in other cancers and has been shown to be expressed in a subset of breast cancers (Noronha et al., 2011; Simon et al., 2004; Tsuda et al., 2005), though c-KIT expression in breast cancer remains unresolved. To our knowledge, there are no previous studies linking c-KIT with endocrine resistance, making this factor an attractive target for further investigation. To follow up on this finding, we first overexpressed c-KIT in MCF-7-S cells and confirmed that these overexpressing cells were more resistant to endocrine therapy. We next investigated how c-KIT might be promoting resistance by looking at potential downstream targets of c-KIT signaling using PRO-seq. Analysis of genes that were upregulated following c-KIT overexpression using the KEGG 2021(Kanehisa, 2019; Kanehisa et al., 2021; Kanehisa & Goto, 2000) and ENRICHR (EY et al., 2013; MV et al., 2016; Xie et al., 2021) databases found that the ABC transporter protein family was enriched in our dataset. Interestingly, ABC transporters have been found to play a role in multi-drug resistance in a range of cancers (Doyle & Ross, 2003; Mohelnikova-Duchonova et al., 2013) and have also been associated with the efflux of TAM and TAM metabolites in breast cancer cells (Domenichini, Adamska, & Falasca, 2019; Hansten, 2018; Teft, Mansell, & Kim, 2011). Moreover, upregulation of ABC transporters has been observed in TAM resistant BC cells (Choi, Yang, Sang, Chang, & Keon, 2007; Dubrovska et al., 2012).

These results help formulate a working hypothesis for how c-KIT signaling could promote TAM resistance (Figure 6E). In our working model, upon upregulation of c-KIT, the c-KIT ligand, stem cell factor (SCF), would bind to c-KIT promoting autophosphorylation and subsequent activation of signaling pathways such as PI3K, SRC, or NRAS (Fig. 3). This signaling would then promote the upregulation of ABC transporter transcripts, which would then be translated and incorporated into the plasma membrane. Activated ABC transporters could then efflux TAM from inside the cell, therefore freeing estrogen receptor alpha (ER*α*) to bind to estrogen (E2) and promote ER*α* downstream signaling, cell proliferation/survival, and TAM resistance. In support of this prediction, the c-KIT inhibitor, Gleevec, has been found to inhibit ABC transporter activity (Chau, Ip, Mak, Lai, & Wong, 2012; Kathawala, Gupta, Ashby, & Chen, 2015; Wu et al., 2019). Highlighting the potential clinical relevance of our findings with respect to ABC transporters, we performed Kaplan Meier plotter analysis of these factors and found that overexpression of ABCA1 or ABCG1 were correlated with a worse prognosis for ER*α*+ BC patients treated with endocrine therapy.

Regarding other potential mechanisms by which c-KIT may promote endocrine resistance, c-KIT has also been found to be important in the maintenance of hematopoietic stem cells (Kimura et al., 2011). In breast tissue, there are distinct populations of progenitor luminal cells that are ER-cKIT+. c-KIT+ luminal cells, but not c-KIT-luminal cells, are progenitors that can be successfully transplanted into mouse fat pads, resulting in mammary epithelial outgrowth and expression of both ER*α*- and ER*α*+ differentiated cells (Regan et al., 2011). Moreover, strong expression of SCF was detected in breast tissue basal cells, suggesting a method to activate c-KIT signaling in the ER-cKIT+ luminal progenitor population. Therefore, it is possible that c-KIT may also be important in the maintenance of BC progenitor cells and potentially important in replenishing tumor cells following endocrine therapy treatment, thereby contributing to BC recurrence and resistance. This phenomenon is observed in non-small lung cancer cells where chemotherapy kills the differentiated cancer cells but not the cancer stem cells (CSCs) (Levina et al., 2010) and in c-KIT+ prostate CSC-like cells which are associated with a decreased overall survival and progression-free survival (Harris et al., 2021). The population of c-KIT+ cells could be restricted to the proportionally smaller cancer stem/progenitor cell population, and the relative scarcity of this population may contribute to the controversy regarding whether or not c-KIT plays an important role in ER*α*+ BC.

With regards to the potential clinical relevance of our work, our inhibitor studies suggest that c-KIT represents a novel target for treating endocrine resistance in women. When we tested for the inhibitory effects of multiple concentrations of TAM and Gleevec treatment on MCF-7-S cell growth, we found there is an additive and/or synergistic inhibitory effect of these drugs on growth at specific concentrations when compared to individual Gleevec and TAM treatments. Importantly, c-KIT inhibitors, such as Gleevec, have already been approved by the FDA to treat c-KIT expressing cancers. Therefore, if our studies are validated by others and c-KIT is found to play a role in endocrine resistance in a clinical setting, then the path towards the use of c-KIT inhibitors in treating endocrine resistance in women may be fairly straightforward.

## Methods

### Cell lines and cell culture

TamS MCF-7 cells were a gift from Dr. Joshua LaBaer (Gonzalez-Malerva et al., 2011). TamS cells containing the dCAS9-VP64-HAtag (addgene # 50916) were used throughout this experiment. Cells were grown in Dulbecco’s Modified Eagle Medium (DMEM) supplemented using a specific lot (Seradigm; Lot # 335B15) of 5% fetal bovine serum (FBS) that tested negative for TET activity and 1X Antibiotic-Antimycotic (Gibco; Cat#15240-062). Phoenix-ampho cells (Allele Biotechnology; cat# ABP-RVC-10001) were grown in Roswell Park Memorial Institute (RPMI)-1640 medium supplemented with 10% FBS and 1X Antibiotic-Antimycotic. The following cell culture media supplements and concentrations were used throughout this study unless indicated otherwise: Doxycycline hyclate (2ug/ml, Sigma, cat# D9891-5G), tamoxifen as (Z)-4-Hydroxytamoxifen (1uM, 4-OHT; Sigma-Aldrich; Cat# H7904), and Gleevec (Imatinib mesylate; Santa Cruz; cat # CAS 220127-57-1).

### Lentivirus production

Phoenix-eco cells were seeded in six 15cm dishes at approximately 9,031,250 cells/dish. After 24hrs, cells were transfected using Mirus Trans-Lenti Transfection Reagent (Cat # MIR 6600) according to the manufacturer’s protocol. Second generation virus particles were made using psPAX (addgene #12260) and pMD2.G (addgene #12259). Lentiviruses were made from the CRISPRa-v2 top 5 sgRNAs/gene pooled plasmid library (addgene # 83978) to develop CRISPRa lentiviruses to modulate target gene transcription. Media containing virus particles was collected after 48hrs, filtered using 0.45um filter, snap frozen, and stored at -80°C until time of infection.

### Lentivirus Transduction

Seventeen million cells were plated per 245mm dish (Corning, cat# 431110) 24hrs prior to the lentivirus transduction. A total of eight dishes were plated per replicate. Two biological replicates were completed. At the time of transduction, there were approximately 29,750,000 cells/dish (238,000,000 cells total). 30ml of virus and media were first added for 2.5hrs and then 70ml of fresh media was added for a total of 100ml. Infected cells were tested using flow cytometry to determine the percentage of BFP positive cells (from the sgRNA plasmids) after 48hrs. Cells were plated in new 245mm dishes and selected with 2ug/ml puromycin (to select for cells containing sgRNA plasmids) for 48hrs or until cells were 80-90% positive (Kampmann et al., 2014). Cells were then plated into four polystyrene CellSTACK 5-layer chambers (Corning, cat# 3319) at 106,000,000 cells per CellSTACK. DOX or vehicle control was added to each chamber. Then, TAM or vehicle control (with DOX or vehicle control) was added to the media after 12-24hrs (Figure 1E). Media was replaced every 48hrs. Cells were split and counted as the top layer of the chamber became confluent. Cells were treated with vehicle or DOX and with vehicle or TAM for 20 days.

### Isolation of genomic DNA

Sequencing libraries were prepared following the “Illumina Sequencing Sample Preparation for use with CRISPRi/a-v2 Libraries” protocol (Weissman, n.d.), with minor modifications. 100M freshly harvested cells were used per treatment. When using the NucleoSpin Blood XL kit (Machery Nagel; cat # 740950.10), the samples were spun at 3700g for 5min instead of the recommended 4000g 3min. The final amount of gDNA obtained for each sample was between 1mg and 1.4mg. In steps 2a and 2b, after digestion, cells were concentrated using 3M sodium acetate and isopropanol and then washed with 75% EtOH. DNA was re-dissolved in 250ul of Milli-Q filtered ultrapure H2O and mixed with 250ul of glycerol loading dye mixture (62.5ul 50x TAE, 312.5ul glycerol, 312.5ul 6x loading dye). Following the protocol mentioned above, samples were run on a 0.8% agarose gel and the regions between 300 to 650bps were cut out to encompass the expected product size of 471bp. Gel extraction was performed using ThermoScientific GeneJET Gel Extraction Kit (Cat #K0692) according to the manufacturer’s specifications. Four to six columns per sample were used. Columns were eluted in 20ul of Milli-Q filtered ultrapure water. After centrifugation, the eluted sample was added to a second column where the incubation was repeated and then centrifuged, resulting in a final volume of 40ul. The Thermo scientific nanodrop 2000/2000c spectrophotometer was used to determine gDNA concentrations.

### qPCR for Illumina Sequencing

qPCR was used to determine the number of cycles needed to amplify the product prior to Illumina sequencing. The qPCR reaction was scaled down to 50ng in 10ul reaction. For each reaction 5ul PowerUp Sybr (Cat# A25741), 1ul of 5uM forward and reverse primer mix, 50ng of sample, and molecular grade water to 10ul was used. The fast cycling mode according to the manufacturer’s instructions was used. The amount of amplification needed to send for sequencing was determined based on the threshold (Ct) after completion of qPCR.

### PCR to add Barcodes and Illumina sequencing

The protocol in “Illumina Sequencing Sample Preparation for use with CRISPRi/a-v2 Libraries” was followed with the following modifications (Weissman, n.d.). The number of cycles were changed depending on the qPCR reaction mentioned above. Two PCR reactions were performed for each sample. Instead of using Phusion, Q5 High-Fidelity 2X Master Mix (NEB #M0492S) was used. Illumina sequencing primers that were added into the PCR reaction are listed in table S3.14. To directly test the PCR reaction an 8% TBE gel and a 2% agarose gel were used instead of a 20% TBE gel. After PCR amplification, 0.8x Ampure beads (Axygen AxyPrep – Cat# MAGZ-PCR-CL-5), following the manufacturers protocol, were used for PCR purification. The 150bp primer dimer was removed and the 274bp product was retained. Samples were prepared for Illumina sequencing according to core facility instructions. NextSeq500 was used for sequencing with the paired end kit. The average sequencing read depth was 40,925,725.

### Differential expression analysis of sgRNAs

A genome index was made of the sgRNA DNA sequence library using bwa (Li & Durbin, 2009) index. sgRNAs from the screen had adapters removed using cutadapt (Martin, 2011), were aligned to the index using bwa aln, and made into bam files using bwa samse and samtools (Li, 2011; Li et al., 2009). Alignments with a mapq score of 20 or greater were then sorted and indexed using samtools. Bedtools multicov (Quinlan & Hall, 2010) was used to count reads in each sgRNA from the original DNA library (giving the number of sgRNAs from the screen in each gene) and convert to a bed file to be used in downstream analyses. R package ‘DESeq2’ (Love, Huber, & Anders, 2014) was used to perform differential expression analysis of the number of sgRNAs in different conditions. Significant differentially expressed genes were defined using the parameters p < 0.05 and log2 fold change ≥ 2.5. Two biological replicates were performed.

### Gene set over-representation analysis

Gene set overrepresentation analysis based on TAM resistant candidate genes (p < 0.05) that were identified using ‘DESeq2’ (Love et al., 2014), was performed by R package ‘HTSanalyzeR2’ (L.Zhu, F. Gao, X. Mei, n.d.) using hypergeometric tests. Significant gene sets were defined by Benjamini-Hochberg adjusted p < 0.05.

### Cell proliferation validation assay

40,000 cells were seeded in a 24-well plate. The next day, cells were treated with 2ug/ml of DOX or vehicle control. 24hrs later, cells were treated with DOX or vehicle control and TAM or vehicle control. Cells were grown until control cells were close to being confluent, 5-9 days. Cells were fixed with 2% PFA, washed with PBS, stained with crystal violet, washed twice with PBS, air dried, and 500ul of 10% acetic acid was added to each well to remove crystal violet stain from cells. Absorbance was read with a Tecan infinite M200PRO plate reader at 592nm. The equation used in analysis is found in Figure 5b.

### qPCR for relative transcription levels

500,000 cells were plated in 6 well plates. 24hrs later, cells were treated with DOX or vehicle control for 24hrs and then RNA was collected and extracted using TRI Reagent (Molecular Research Center, Inc., cat # TR118). cDNA was made using High-Capacity RNA-to-cDNA kit (Applied Biosystems, cat# 4387406) by following the manufacturer’s instructions. DNA was diluted 1:5 before using for qPCR. PowerUp SYBR Green was used according to the manufacturer’s instructions (Life Technologies Cat#A25741). Primers used are found in Table S14.

### Cell culture and nuclei isolation

5,000,000 cells were seeded in 150mm TC-treated culture dishes. 24hrs later cells were treated with DOX or vehicle control. The second experiment (Figure 6c) was then treated with DOX or vehicle control and with TAM for 5 days. Cells were rinsed with PBS, trypsinized, quenched with media, and centrifuged for 5 minutes at 4°C. Media was aspirated off, cells were washed in 1ml cold PBS, twice. Cells were then spun and resuspended in 150ul wash buffer (1ml of stock solution [10mM Tris-HCl pH8, 2mM MgAc, 10mM NaCl, 300mM sucrose], 0.5mM DTT, Protease Inhibitor Cocktail EDTA-free (ThermoFisher; cat# A32955), and 20 units/ml SUPERase In RNase Inhibitor (Life Technologies; AM2694)). 150ul of 2X lysis buffer (1ml of stock [10mM Tris-HCl pH8, 2mM MgAc, 10mM NaCl, 300mM sucrose, 6 mM CaCl2, 0.2% NP-40], 0.5mM DTT, Protease Inhibitor Cocktail EDTA-free, and 20 units/ml SUPERase In RNase Inhibitor. Cells were then pipetted up and down 10 times to facilitate nuclei release then centrifuged at 1000g for 5 min at 4°C. Supernatant was aspirated off, cells were resuspended in 500ul wash buffer and 500ul 2X lysis buffer, and centrifuged at 1000 x g for 5 min at 4°C. Supernatant was aspirated off, nuclei was resuspended in 50ul storage buffer (1ml of stock [50mM Tris-HCl pH 8.3, 40% glycerol, 5mM MgCl2, 0.1 mM EDTA, pH 8.0], plus 4 units/ml SUPERase In RNase Inhibitor). Nuclei were stained and counted using trypan blue.

### Nuclear run-on and PRO-seq library preparation

Nuclear run-on experiments were performed according to previously described methods (Kwak, Fuda, Core, & Lis, 2013; Mahat et al., 2016). Nuclei stored in 50ul freezing buffer were thawed on ice. Once thawed, nuclei were mixed with 50ul of equilibrated (to 37°C) 2X biotin run-on master mix (10 mM Tris-HCl pH 8.0, 5mM MgCl2, 1mM DTT, 300mM KCl, 400μM ATP (Thermo Scientific; Cat# R0441), 400μM GTP (Thermo Scientific; Cat# R0461), 40μM biotin-11-CTP (Perkin Elmer Cat# NEL542001), 40μM biotin-11-UTP (Perkin Elmer; Cat# NEL543001), 0.8 units/μL SUPERase Inhibitor (Invitrogen; Cat# AM2696), 1% Sarkosyl). The mixture was incubated for 5 min at 37°C. The run-on reaction was stopped using 500ul of TRI reagent LS (Molecular Research Center; cat# TS 120) and incubated at room temperature for 5 minutes. RNA was then isolated following the manufacturer’s instructions and GlycoBlue (Life Technologies; cat# AM9515) was used to identify the RNA pellet and obtain a higher yield. RNA pellets were re-dissolved in 20ul DEPC water at 4°C, heated at 65°C for 40 seconds to denature secondary structures, then immediately placed on ice. 5ul of 1N NaOH was added to each sample and incubated on ice for 4min. 25ul of 1M Tris-HCl pH 6.8 was added to each sample to neutralize the base hydrolysis reaction. Bio-Rad P-30 columns (cat# 732-6250) were prepared and used following manufacturer’s instructions. RNA from columns were eluted in 10ul DEPC water for magnetic bead binding. After RNA binding to magnetic beads, beads were washed using salt washes, RNA was removed from beads using TRI reagent (Molecular Research Center; cat# TS 118), and 3’ RNA adaptor ligation was completed following methods referenced above. The following day, magnetic bead binding, salt washes, and TRI reagent for RNA extraction was again performed followed by 5’ de-capping, hydroxyl repair, and 5’ RNA adaptor ligation. Bead binding was again performed as mentioned above, followed by reverse transcription (SuperScript IV Reverse Transcriptase (Life Technologies; cat# 18090050). Library amplification was performed as described in qPCR for Illumina Sequencing methods section from above and samples were run on 8% TBE gel, extracted and then eluted to remove primer dimers as described in the methods referenced above. Samples were further prepared according to the Cornell University Institute of Biotechnology Sequencing Core instructions and sequenced using an Illumina NextSeq500.

### Differential expression analysis of PRO-seq data

PRO-seq data was mapped to hg38using the methods described in (Kwak et al., 2013; Mahat et al., 2016). Briefly, PRO-seq data was trimmed using cutadapt and mapped to hg38 using bwa. BedTools and bedGraphToBigWig were used to convert aligned reads to the bigwig format. To compare treatment conditions, differential expression analysis was conducted using DESeq2(Love et al., 2014) with GENCODE_v38 gene annotations. Differentially upregulated expressed genes between conditions were defined using an adjusted p-value (Benjamini-Hochberg) of less than 0.075.

### Drug combination synergy

15,000 TamS cells were seeded in 48 well plates. After 24hrs each well was treated with vehicle control, TAM, Gleevec, or both at increasing concentrations. Cells were grown for 5 days or until confluent. Two technical replicates were performed for each experiment and three biological replicates were performed. Cells were fixed as described in the cell proliferation validation assay section above. Synergism was determined using SynergyFinder 2.0(Ianevski et al., 2020). The ZIP method was used, and synergism was determined using the parameters described in the SynergyFinder user guide.

### Protein analysis

Cells were seeded at 500,000 cells per plate, treated with DOX or vehicle control, and collected after 24hrs. Cells were washed with ice-cold PBS before lysing with radioimmunoprecipitation assay buffer (RIPA) buffer and Pierce protease inhibitor (Thermo Scientific, ref# A32955). Cells were removed using a cell lifter over ice and spun down at 13,000g at 4°C for 20 minutes. Supernatant was stored at -20°C. A micro bicinchoninic acid (BCA) assay (Thermo Scientific, cat # 23235) was used according to the manufacturer’s instructions to determine protein concentration. A Bio-Rad Trans-Blot Turbo western blot semi-dry transfer system was used for all transfers using their proprietary transfer buffer and stacks. Transfer was up to 25V and 1.3 amps for 10 minutes. Protein was transferred to 0.2um PVDF membrane (IMMUNOBILON-P^SQ^ cat # ISEQ00010). Membranes were blocked in 5% non-fat milk for 1hr at room temperature. All antibodies were diluted in 5% non-fat milk and incubated overnight at 4°C. Anti-HA tag (F-7) (Santa Cruz; cat # sc-7392) and B-Actin (AbCam, ab8227) antibodies were used at 1:500 and 1:5000, respectively. Membranes were washed in tris buffered saline with 1% tween (TBST). Secondary anti-mouse and anti-rabbit peroxidase-conjugated affiniPure antibodies from Jackson ImmunoResearch Laboratories (cat# 115-035-146 and cat #111-035-144, respectively) were diluted to 1:10000 and 1:20000, respectively, in TBST and incubated at room temperature for 1hr. Membranes were washed 3X for 20 minutes with TSBT and then imaged using WesternBright Quantum detection kit (cat# K-12042-D10) and a Bio-Rad ChemiDoc MP.

### Statistics

Student’s one-tailed T-Tests were used in Figure 4 and two-tailed T-Tests were used in Figure 5. Samples were normalized to controls prior to analysis. Three biological replicates were used, unless otherwise specified.

---

**S1 Fig.**
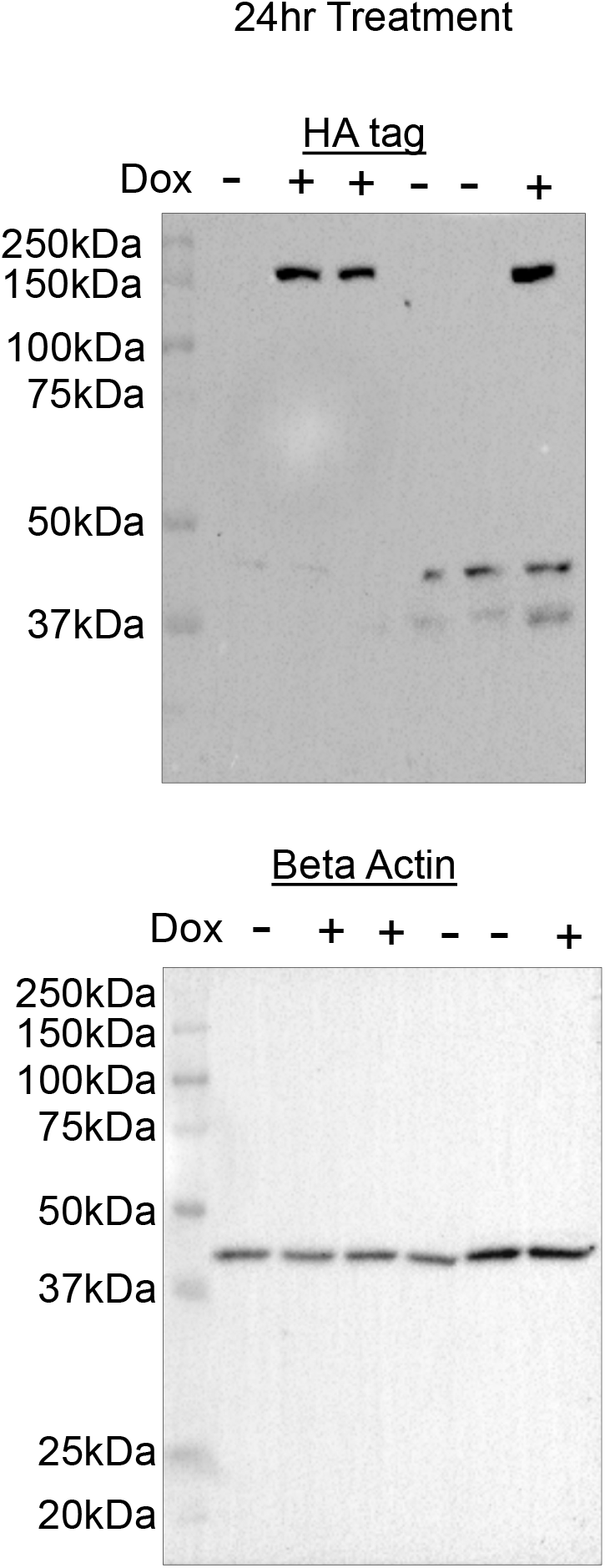
Western blot of HA-dCAS9-VP64 fusion protein and b-Actin expression following 24hr vehicle or DOX treatment.

**S2 Fig.**
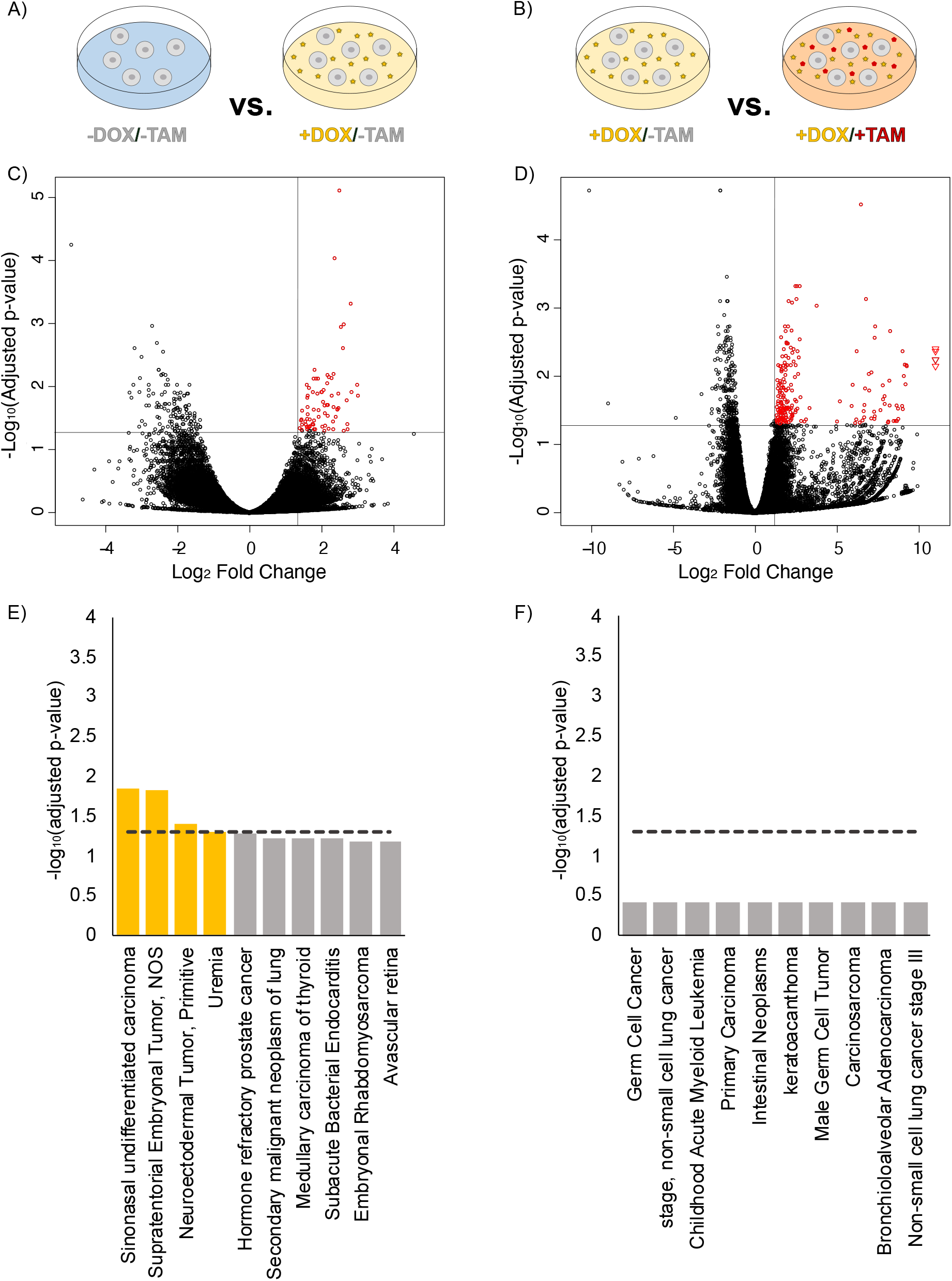
Analysis of CRISPRa screens in the non-experimental groups did not identify any genes associated with BC or resistance. A-B) Schematic representation of treatment conditions corresponding to the volcano plots and bar charts shown below them. C) Volcano plot in which the upper right quadrant identifies sgRNAs (genes) that were significantly over-represented when comparing cells with CRISPR induction and without TAM treatment (+DOX/-TAM) to cells in the absence of CRISPR induction or TAM treatment (-DOX/-TAM), (p < 0.05). D) Volcano plot in which the upper right quadrant identifies sgRNAs (genes) that were significantly over-represented when comparing cells after CRISPR induction and TAM treatment (+DOX/+TAM) to cells after CRISPR induction and without TAM (+DOX/-TAM), (p < 0.05). Circles represent individual sgRNAs, and outliers are represented as triangles. E-F) DISGENET disease enrichment database (within the ENRICHR database) was used to identify the top 10 human diseases associated with the treatment specific subsets of enriched sgRNAs (genes) identified in the volcano plots above, showing that human diseases in the non-experimental comparisons have not been previously implicated in breast cancer or resistance. Dotted line represents the adjusted p-value < 0.05. *NOS = Not Otherwise Specified.

**Table S1.** Read Counts.

| Dox Treatment | TAM treatment | Replicate | Read Counts |
| --- | --- | --- | --- |
| No | No | 1 | 71,819,615 |
| No | Yes | 1 | 31,110,019 |
| Yes | No | 1 | 84,133,329 |
| Yes | Yes | 1 | 28,095,276 |
| No | No | 2 | 31,156,609 |
| No | Yes | 2 | 29,598,316 |
| Yes | No | 2 | 19,982,851 |
| Yes | Yes | 2 | 31,509,781 |
|  |  | average | 40,925,725 |

**Table S2.** Significantly enriched gene comparison list for treatment comparison -Dox/-Tam vs. – Dox/+Tam.

| **names** | **indx** | **padj_1.3 (0.05)** | **log2foldchange_1.32 (2.5)** |
| --- | --- | --- | --- |
| **ACTR1A** | 1208 | 5.12E-03 | 10.46 |
| **ALDH4A1** | 2845 | 8.65E-04 | 7.82 |
| **ALS2CR12** | 3102 | 5.33E-03 | 2.42 |
| **ANKRD10** | 3557 | 5.74E-03 | 7.13 |
| **APBB2** | 4286 | 4.01E-02 | 2.43 |
| **ARHGEF12** | 5086 | 3.16E-02 | 8.94 |
| **ARMC12** | 5440 | 3.87E-02 | 7.59 |
| **ATMIN** | 6367 | 4.76E-02 | 9.42 |
| **BBC3** | 7566 | 3.16E-02 | 8.55 |
| **BGN** | 8062 | 2.40E-02 | 7.82 |
| **BLVRA** | 8269 | 1.26E-02 | 1.81 |
| **C10orf113** | 9114 | 3.09E-02 | 4.38 |
| **C12orf55** | 9533 | 3.75E-02 | 1.61 |
| **C16orf89** | 9958 | 9.04E-03 | 10.71 |
| **C1orf216** | 10752 | 8.66E-03 | 1.79 |
| **C4orf33** | 11538 | 2.81E-08 | 23.28 |
| **C9orf47** | 12259 | 2.46E-02 | 7.74 |
| **CACNA1A** | 12504 | 4.71E-02 | 2.47 |
| **CCDC121** | 13741 | 1.14E-07 | 23.09 |
| **CCNT2** | 14771 | 7.90E-03 | 1.60 |
| **CD3EAP** | 15198 | 4.22E-02 | 9.48 |
| **CIAPIN1** | 17536 | 4.08E-02 | 1.57 |
| **CIZ1** | 17673 | 3.75E-02 | 9.56 |
| **CLEC12B** | 18018 | 1.48E-06 | 22.86 |
| **CLIC5** | 18197 | 1.00E-02 | 7.48 |
| **CNTD2** | 18858 | 2.76E-02 | 1.43 |
| **COG2** | 18999 | 5.01E-02 | 7.96 |
| **COPS8** | 19467 | 4.04E-02 | 1.35 |
| **CTNNBIP1** | 21008 | 3.81E-03 | 8.02 |
| **CUEDC1** | 21197 | 9.33E-03 | 7.83 |
| **DBT** | 22301 | 7.90E-03 | 9.99 |
| **DEFB124** | 23169 | 3.75E-02 | 8.88 |
| **DGCR6** | 23468 | 6.42E-04 | 2.79 |
| **DLG3** | 23957 | 1.73E-03 | 2.11 |
| **DLK1** | 24012 | 3.87E-02 | 1.83 |
| **DSC1** | 25162 | 4.07E-02 | 1.90 |
| **DUOXA1** | 25368 | 3.70E-02 | 1.70 |
| **DYNLT1** | 25646 | 5.74E-03 | 2.80 |
| **EIF2B1** | 26485 | 1.66E-06 | 21.29 |
| **EIF3G** | 26561 | 2.26E-02 | 7.12 |
| **EIF5** | 26700 | 8.73E-03 | 6.35 |
| **FAM101B** | 28786 | 2.20E-03 | 10.59 |
| **FAM102B** | 28799 | 4.68E-02 | 5.85 |
| **FAM167A** | 29295 | 3.48E-02 | 8.28 |
| **FAM3A** | 29933 | 4.26E-02 | 7.30 |
| **FBXO9** | 30936 | 4.19E-02 | 1.91 |
| **FOXA3** | 32173 | 1.01E-02 | 7.15 |
| **GATA3** | 33644 | 2.20E-02 | 2.10 |
| **GBP4** | 33766 | 1.71E-02 | 1.42 |
| **GEM** | 34098 | 4.16E-02 | 2.33 |
| **GFAP** | 34154 | 5.01E-02 | 7.26 |
| **GIGYF2** | 34402 | 4.27E-02 | 1.81 |
| **GKAP1** | 34669 | 3.34E-02 | 1.84 |
| **GRP** | 36734 | 1.93E-02 | 7.14 |
| **JAZF1** | 43289 | 2.77E-03 | 1.65 |
| **KBTBD2** | 43580 | 4.35E-02 | 7.67 |
| **KLHL36** | 45547 | 1.48E-03 | 2.43 |
| **KRTAP10-8** | 46317 | 2.25E-03 | 2.00 |
| **LANCL3** | 46994 | 4.15E-02 | 3.68 |
| **LEPREL2** | 47542 | 1.26E-02 | 9.23 |
| **LPPR2** | 48938 | 3.14E-02 | 8.42 |
| **MAST3** | 51322 | 1.93E-02 | 1.82 |
| **MGAT4B** | 52634 | 2.71E-02 | 7.26 |
| **MYH15** | 55473 | 4.68E-02 | 4.29 |
| **NHSL1** | 57694 | 5.46E-03 | 10.09 |
| **NLGN4X** | 58000 | 5.77E-03 | 5.27 |
| **NPPA** | 58735 | 3.67E-02 | 1.67 |
| **NPR 1.00** | 58747 | 3.09E-02 | 7.37 |
| **NSUN6** | 59271 | 3.14E-02 | 1.53 |
| **OR1Q1** | 60876 | 3.09E-02 | 8.06 |
| **OR7A10** | 62093 | 1.12E-02 | 2.63 |
| **PARS2** | 63461 | 2.81E-08 | 2.86 |
| **PCED1B** | 64069 | 5.01E-02 | 1.99 |
| **PDE4DIP** | 64466 | 5.78E-03 | 7.31 |
| **PKIA** | 66327 | 3.16E-02 | 8.17 |
| **PKIB** | 66331 | 2.12E-03 | 2.03 |
| **PKN2** | 66376 | 9.50E-03 | 7.99 |
| **PLA2G4D** | 66508 | 3.39E-03 | 6.94 |
| **POSTN** | 67976 | 8.66E-03 | 1.77 |
| **PRR4** | 69917 | 7.71E-03 | 1.86 |
| **PTTG1IP** | 71266 | 3.16E-02 | 1.61 |
| **RB1CC1** | 72784 | 1.76E-02 | 2.10 |
| **RNF135** | 74760 | 2.86E-02 | 6.72 |
| **RPGRIP1** | 75358 | 4.38E-02 | 1.36 |
| **SCMH1** | 77410 | 3.34E-02 | 1.64 |
| **SLC28A2** | 80864 | 5.46E-03 | 1.91 |
| **SLC39A10** | 81311 | 1.48E-02 | 1.76 |
| **SMAD6** | 82204 | 1.14E-07 | 23.11 |
| **SMIM18** | 82477 | 1.10E-02 | 1.74 |
| **SOD3** | 83196 | 5.57E-03 | 3.43 |
| **SSTR4** | 84942 | 7.43E-03 | 8.09 |
| **SUCLG1** | 85829 | 1.69E-02 | 2.55 |
| **SUSD3** | 86128 | 2.52E-02 | 7.69 |
| **TBC1D13** | 87264 | 4.22E-02 | 5.77 |
| **TEF** | 88126 | 4.26E-02 | 6.33 |
| **TMC4** | 89635 | 5.45E-03 | 4.37 |
| **TMEM88B** | 91010 | 1.05E-05 | 2.49 |
| **TNFRSF10A** | 91357 | 4.34E-03 | 1.66 |
| **TNFRSF17** | 91415 | 3.75E-02 | 6.67 |
| **TNN** | 91608 | 1.43E-02 | 7.22 |
| **TOR1AIP2** | 91896 | 3.44E-02 | 1.72 |
| **TOX2** | 91928 | 7.52E-06 | 7.49 |
| **TPBG** | 92022 | 6.82E-03 | 7.79 |
| **TRIM27** | 92668 | 2.77E-03 | 2.77 |
| **TSPAN10** | 93485 | 3.05E-02 | 2.64 |
| **UCK1** | 95131 | 4.37E-02 | 7.35 |
| **UTP23** | 96128 | 3.39E-03 | 5.46 |
| **VSTM2B** | 96831 | 3.14E-02 | 9.66 |
| **ZC3HAV1L** | 98799 | 5.01E-02 | 1.63 |
| **ZNF268** | 100082 | 3.34E-02 | 7.91 |
| **ZNF490** | 100792 | 2.42E-02 | 6.97 |
| **ZNF93** | 102296 | 4.00E-02 | 2.42 |
| **ZNRF3** | 102353 | 8.66E-03 | 7.61 |
| **ZRSR2** | 102416 | 1.61E-02 | 2.32 |
| **ZSWIM6** | 102571 | 4.19E-02 | 7.67 |
| **non-targeting** | 102969 | 9.78E-08 | 23.15 |
| **non-targeting** | 103634 | 2.52E-02 | 9.75 |
| **non-targeting** | 103695 | 2.20E-03 | 3.87 |

**Table S3.** Significantly enriched gene comparison list for treatment comparison -Dox/-Tam vs. +Dox/-Tam.

| **names** | **indx** | **padj_1.3 (0.05)** | **log2foldchange_1.32 (2.5)** |
| --- | --- | --- | --- |
| **A4GALT** | 31 | 1.28E-02 | 1.89 |
| **AGER** | 2146 | 1.98E-02 | 2.09 |
| **ALS2CR12** | 3102 | 2.29E-02 | 2.45 |
| **ANKRD28** | 3667 | 2.34E-02 | 1.92 |
| **ANXA7** | 4054 | 2.61E-02 | 2.10 |
| **APOBEC3B** | 4459 | 5.00E-02 | 2.09 |
| **ARID4A** | 5244 | 3.31E-02 | 1.45 |
| **ARTN** | 5736 | 1.39E-02 | 2.29 |
| **ASCL1** | 5918 | 9.03E-05 | 2.37 |
| **B4GALT5** | 7286 | 3.89E-02 | 1.76 |
| **BHLHE41** | 8099 | 3.02E-02 | 1.93 |
| **C9orf69** | 12289 | 4.14E-02 | 2.11 |
| **CALCB** | 12728 | 1.74E-02 | 2.05 |
| **CCRL2** | 14856 | 1.39E-02 | 3.02 |
| **CDHR4** | 15780 | 2.26E-02 | 2.45 |
| **CIB2** | 17549 | 3.02E-02 | 2.40 |
| **CTR9** | 21043 | 1.39E-02 | 1.62 |
| **DLC1** | 23926 | 1.21E-02 | 1.77 |
| **DPY19L2** | 25026 | 4.70E-02 | 1.83 |
| **EGFR** | 26308 | 7.50E-03 | 1.91 |
| **ENPP7** | 27301 | 1.28E-02 | 2.47 |
| **F13B** | 28566 | 2.08E-02 | 1.62 |
| **FAM46D** | 29987 | 3.97E-02 | 1.78 |
| **FAR2** | 30452 | 6.57E-03 | 2.17 |
| **FERMT1** | 31260 | 9.35E-03 | 3.00 |
| **FTCD** | 32729 | 1.96E-02 | 2.18 |
| **GJC1** | 34613 | 2.42E-02 | 1.47 |
| **GLCCI1** | 34714 | 8.75E-03 | 2.17 |
| **GPNMB** | 35743 | 2.20E-02 | 2.48 |
| **GRAP2** | 36438 | 4.81E-04 | 2.82 |
| **GRB2** | 36474 | 4.52E-02 | 1.40 |
| **GSTM3** | 36928 | 3.97E-02 | 1.57 |
| **HIST1H2BG** | 38642 | 4.70E-02 | 1.80 |
| **HSD17B10** | 40042 | 7.50E-03 | 1.79 |
| **IL15** | 41523 | 4.31E-02 | 1.61 |
| **IMMT** | 41997 | 5.36E-03 | 1.81 |
| **IQGAP2** | 42536 | 2.11E-02 | 2.34 |
| **KDR** | 44368 | 1.05E-02 | 1.66 |
| **KRTAP5-9** | 46633 | 5.00E-02 | 2.62 |
| **KXD1** | 46747 | 4.42E-02 | 1.65 |
| **LINGO3** | 48064 | 4.70E-02 | 2.73 |
| **LRP3** | 49120 | 4.78E-02 | 1.46 |
| **MDK** | 51830 | 3.54E-02 | 1.55 |
| **MRPS9** | 54342 | 3.42E-02 | 1.44 |
| **NGEF** | 57614 | 4.39E-02 | 1.61 |
| **NUTM1** | 59812 | 9.35E-03 | 1.60 |
| **OR6C3** | 61983 | 4.77E-02 | 1.61 |
| **OR8B12** | 62142 | 1.64E-02 | 2.71 |
| **OR8B4** | 62160 | 4.24E-02 | 1.68 |
| **OTX1** | 62640 | 1.21E-02 | 2.02 |
| **PLAG1** | 66580 | 1.21E-02 | 2.84 |
| **PLCB2** | 66635 | 1.03E-03 | 2.62 |
| **PPP6R3** | 68895 | 1.12E-03 | 2.55 |
| **PTCHD3** | 70716 | 7.62E-06 | 2.50 |
| **RAPGEF4** | 72479 | 1.21E-02 | 2.01 |
| **RARG** | 72524 | 1.96E-02 | 2.16 |
| **RBP7** | 73184 | 3.26E-02 | 1.61 |
| **RUFY4** | 76449 | 1.13E-02 | 2.07 |
| **SAYSD1** | 77120 | 3.58E-02 | 2.33 |
| **SERPINF1** | 78580 | 4.70E-02 | 2.34 |
| **SETMAR** | 78727 | 2.29E-02 | 1.79 |
| **SIMC1** | 79682 | 3.95E-02 | 1.47 |
| **SMCO4** | 82373 | 4.31E-02 | 1.68 |
| **SMNDC1** | 82566 | 4.63E-02 | 2.20 |
| **SOX21** | 83389 | 3.42E-02 | 1.80 |
| **SP8** | 83515 | 7.34E-03 | 2.29 |
| **SPINK13** | 84053 | 4.31E-02 | 2.03 |
| **SPRY4** | 84348 | 7.50E-03 | 1.86 |
| **SRXN1** | 84798 | 2.75E-02 | 1.63 |
| **TMEM14C** | 90044 | 6.22E-03 | 2.34 |
| **TMEM180** | 90222 | 3.27E-02 | 1.68 |
| **TMEM248** | 90613 | 2.87E-02 | 2.18 |
| **TNS4** | 91743 | 7.02E-03 | 2.21 |
| **WIPF3** | 97595 | 3.88E-02 | 2.72 |
| **ZRSR2** | 102416 | 2.43E-03 | 2.60 |
| **non-targeting** | 102976 | 1.49E-02 | 1.61 |

**Table S4.** Significantly enriched gene comparison list for treatment comparison +Dox/-Tam vs. +Dox/+Tam.

| **names** | **indx** | **padj_1.3 (0.05)** | **log2foldchange_1.32 (2.5)** |
| --- | --- | --- | --- |
| **A3GALT2** | 30 | 3.31E-02 | 1.51 |
| **ACTL6B** | 1156 | 4.49E-02 | 8.69 |
| **ADAMTS16** | 1495 | 1.12E-02 | 1.38 |
| **ADC** | 1631 | 1.31E-02 | 5.98 |
| **AGBL5** | 2143 | 4.92E-02 | 1.56 |
| **ALDOB** | 2880 | 4.48E-02 | 1.35 |
| **AMY2A** | 3319 | 4.68E-02 | 1.75 |
| **ANAPC11** | 3350 | 4.23E-02 | 1.32 |
| **ANKEF1** | 3490 | 4.23E-02 | 6.20 |
| **ANKFN1** | 3494 | 1.85E-03 | 2.04 |
| **ANKRD10** | 3557 | 2.15E-02 | 7.83 |
| **AQPEP** | 4710 | 3.14E-02 | 1.74 |
| **ARHGEF15** | 5095 | 3.36E-02 | 1.49 |
| **ARRDC2** | 5634 | 8.43E-03 | 1.64 |
| **ASIC4** | 5989 | 2.93E-02 | 8.75 |
| **ATF2** | 6203 | 3.70E-02 | 2.30 |
| **ATF7IP** | 6245 | 4.17E-02 | 1.77 |
| **ATP6AP1L** | 6682 | 9.14E-03 | 6.96 |
| **ATP6V1E1** | 6774 | 4.39E-02 | 1.39 |
| **ATP6V1G3** | 6800 | 4.50E-02 | 1.83 |
| **AUNIP** | 6980 | 1.76E-02 | 1.67 |
| **BAG1** | 7358 | 2.84E-02 | 2.05 |
| **BRD3** | 8640 | 4.43E-02 | 2.19 |
| **C12orf10** | 9453 | 4.74E-02 | 8.14 |
| **C17orf104** | 10017 | 2.69E-02 | 1.76 |
| **C17orf51** | 10063 | 2.93E-02 | 2.61 |
| **C2orf54** | 11210 | 2.80E-02 | 1.35 |
| **C5orf63** | 11726 | 3.01E-02 | 2.15 |
| **C9orf47** | 12259 | 2.31E-02 | 5.20 |
| **CAMKK2** | 12907 | 3.96E-03 | 1.77 |
| **CAMTA2** | 12951 | 9.14E-04 | 3.78 |
| **CBFA2T3** | 13458 | 1.14E-02 | 8.22 |
| **CCDC125** | 13762 | 3.29E-02 | 1.49 |
| **CCDC148** | 13871 | 2.43E-02 | 1.61 |
| **CCDC84** | 14336 | 4.47E-02 | 1.85 |
| **CCND1** | 14666 | 7.35E-04 | 2.55 |
| **CCND1** | 14669 | 4.69E-02 | 1.94 |
| **CD69** | 15297 | 4.29E-03 | 22.48 |
| **CEBPD** | 16240 | 3.02E-02 | 9.05 |
| **CEP128** | 16493 | 2.93E-02 | 1.62 |
| **CITED1** | 17655 | 5.11E-03 | 1.69 |
| **CLIC5** | 18197 | 2.73E-03 | 7.32 |
| **CLNS1A** | 18302 | 1.26E-02 | 1.55 |
| **CNN1** | 18708 | 2.93E-02 | 1.97 |
| **CTDSPL** | 20944 | 6.86E-03 | 9.26 |
| **CWH43** | 21306 | 1.50E-02 | 1.41 |
| **CXCL5** | 21380 | 8.00E-03 | 1.64 |
| **CYP8B1** | 21984 | 3.01E-02 | 1.97 |
| **DCDC1** | 22444 | 4.62E-02 | 2.07 |
| **DMKN** | 24104 | 2.90E-02 | 1.54 |
| **DUSP19** | 25455 | 1.62E-02 | 2.28 |
| **DUSP3** | 25493 | 5.73E-03 | 21.95 |
| **DYRK2** | 25667 | 1.62E-02 | 6.99 |
| **EBF4** | 25813 | 2.16E-03 | 8.27 |
| **EGFR** | 26307 | 4.76E-04 | 2.58 |
| **EIF4G2** | 26675 | 2.68E-02 | 8.67 |
| **EIF5** | 26700 | 1.36E-02 | 7.82 |
| **EXOC7** | 28388 | 3.96E-03 | 2.37 |
| **FAM120C** | 28955 | 2.00E-02 | 1.57 |
| **FAM127C** | 29005 | 1.96E-02 | 7.93 |
| **FAM154B** | 29164 | 8.31E-04 | 2.28 |
| **FAM186B** | 29472 | 4.48E-02 | 1.46 |
| **FAM3A** | 29933 | 3.22E-02 | 6.65 |
| **FAS** | 30484 | 3.36E-02 | 1.60 |
| **FCN1** | 31105 | 4.82E-02 | 6.13 |
| **FGF11** | 31390 | 2.60E-02 | 6.70 |
| **FGF4** | 31460 | 1.02E-02 | 2.45 |
| **FNDC7** | 32057 | 1.46E-02 | 1.75 |
| **FOXH1** | 32241 | 2.66E-02 | 8.24 |
| **FZR1** | 33034 | 2.87E-02 | 1.97 |
| **GABRA6** | 33174 | 2.91E-02 | 2.24 |
| **GAD2** | 33246 | 6.78E-03 | 1.44 |
| **GATA1** | 33626 | 4.99E-03 | 1.90 |
| **GATA2** | 33639 | 2.89E-02 | 1.72 |
| **GPR176** | 36015 | 2.21E-02 | 2.46 |
| **GRPEL1** | 36736 | 3.97E-02 | 6.68 |
| **HAPLN1** | 37544 | 4.52E-02 | 8.60 |
| **HEPH** | 38129 | 2.93E-02 | 1.93 |
| **HEPHL1** | 38132 | 1.85E-03 | 7.35 |
| **HEPHL1** | 38134 | 9.14E-03 | 2.65 |
| **HHEX** | 38331 | 3.33E-02 | 2.18 |
| **HHEX** | 38333 | 4.02E-02 | 1.75 |
| **HSF4** | 40149 | 9.78E-03 | 9.11 |
| **HSPBAP1** | 40321 | 1.64E-02 | 1.54 |
| **HTRA3** | 40490 | 4.89E-02 | 1.55 |
| **IFNLR1** | 41041 | 1.85E-03 | 1.85 |
| **IL17RE** | 41591 | 4.20E-02 | 8.77 |
| **IL20RB** | 41722 | 3.39E-02 | 1.86 |
| **INSM1** | 42288 | 4.17E-02 | 7.30 |
| **IQCF6** | 42496 | 1.57E-02 | 1.89 |
| **IRG1** | 42684 | 4.52E-02 | 2.83 |
| **ITGAM** | 42970 | 3.75E-02 | 2.31 |
| **KAT2A** | 43487 | 3.91E-02 | 2.23 |
| **KCNN2** | 44045 | 2.47E-02 | 1.63 |
| **KIDINS220** | 44927 | 2.01E-02 | 2.79 |
| **KIF2C** | 45080 | 4.56E-02 | 8.16 |
| **KIR2DL4** | 45168 | 1.60E-02 | 1.92 |
| **KIT** | 45216 | 7.86E-04 | 2.07 |
| **KRTAP4-7** | 46569 | 2.03E-02 | 1.74 |
| **KSR1** | 46724 | 3.17E-03 | 1.93 |
| **L2HGDH** | 46783 | 2.11E-03 | 2.25 |
| **LETMD1** | 47576 | 9.53E-03 | 1.87 |
| **LMNB2** | 48268 | 3.36E-02 | 1.63 |
| **MAGEA4** | 50363 | 8.72E-03 | 7.13 |
| **MAP4K1** | 50928 | 6.75E-03 | 9.16 |
| **MGAT4B** | 52634 | 3.84E-02 | 7.33 |
| **MUSK** | 55211 | 2.68E-02 | 2.28 |
| **NEDD4L** | 57110 | 3.30E-02 | 7.29 |
| **NIPA1** | 57764 | 2.40E-02 | 1.96 |
| **OPN4** | 60426 | 5.39E-03 | 2.06 |
| **OR11H4** | 60655 | 3.31E-03 | 2.02 |
| **OR51G2** | 61515 | 3.74E-03 | 2.20 |
| **OR51I1** | 61518 | 3.75E-02 | 1.59 |
| **OR6K6** | 62037 | 7.12E-03 | 1.89 |
| **OSBPL5** | 62417 | 3.03E-02 | 1.58 |
| **OVOL1** | 62675 | 4.76E-04 | 2.49 |
| **OXER1** | 62704 | 7.12E-03 | 1.54 |
| **P2RX4** | 62767 | 2.72E-02 | 5.92 |
| **P2RY12** | 62801 | 2.87E-03 | 2.75 |
| **P2RY13** | 62810 | 1.50E-02 | 2.06 |
| **P2RY4** | 62821 | 3.01E-02 | 2.15 |
| **PALM3** | 63146 | 3.96E-03 | 22.67 |
| **PATE3** | 63503 | 4.76E-04 | 2.75 |
| **PCDHB15** | 63895 | 4.50E-02 | 1.86 |
| **PDE6D** | 64494 | 7.35E-04 | 6.79 |
| **PDIA5** | 64617 | 1.31E-02 | 9.21 |
| **PEX6** | 65060 | 4.23E-02 | 3.14 |
| **PGGT1B** | 65272 | 4.26E-02 | 2.02 |
| **PIN4** | 66096 | 6.78E-03 | 2.20 |
| **PIWIL1** | 66246 | 3.62E-02 | 1.58 |
| **PKIB** | 66335 | 1.18E-02 | 1.55 |
| **PLEKHB1** | 66826 | 2.93E-02 | 1.59 |
| **PLEKHF2** | 66850 | 2.16E-03 | 1.89 |
| **POF1B** | 67531 | 7.00E-03 | 1.75 |
| **PORCN** | 67970 | 4.55E-02 | 3.03 |
| **PPARGC1B** | 68231 | 2.33E-02 | 1.43 |
| **PPP5C** | 68869 | 3.01E-05 | 6.49 |
| **PRB1** | 69039 | 4.53E-02 | 1.59 |
| **PRELP** | 69214 | 2.69E-02 | 9.00 |
| **PRKAR1B** | 69368 | 5.73E-03 | 21.95 |
| **PRKCE** | 69404 | 3.39E-02 | 1.79 |
| **PRR15** | 69850 | 1.94E-02 | 1.43 |
| **PRSS42** | 70119 | 4.09E-02 | 7.12 |
| **PSMB7** | 70422 | 8.00E-03 | 1.65 |
| **PURA** | 71307 | 6.78E-03 | 2.50 |
| **RAB3IL1** | 71905 | 1.31E-02 | 1.63 |
| **RAP1B** | 72425 | 4.50E-02 | 1.64 |
| **RASSF9** | 72746 | 2.50E-02 | 1.42 |
| **RBM48** | 73069 | 7.12E-03 | 9.29 |
| **RNF166** | 74858 | 3.31E-03 | 1.95 |
| **RNMTL1** | 75155 | 1.10E-02 | 1.76 |
| **RPL38** | 75593 | 2.55E-02 | 7.94 |
| **RPRM** | 75735 | 4.73E-02 | 6.41 |
| **SAMD12** | 76891 | 1.72E-02 | 1.54 |
| **SAMD3** | 76917 | 1.46E-02 | 1.65 |
| **SCAP** | 77241 | 3.33E-02 | 1.60 |
| **SDR9C7** | 77777 | 4.29E-03 | 2.61 |
| **SFTPC** | 78921 | 3.62E-02 | 1.90 |
| **SLC11A1** | 79994 | 1.20E-02 | 1.82 |
| **SLC26A11** | 80777 | 2.34E-02 | 1.62 |
| **SLC38A7** | 81293 | 4.47E-03 | 2.29 |
| **SLC6A16** | 81711 | 4.93E-02 | 1.53 |
| **SOX21** | 83386 | 6.22E-03 | 1.83 |
| **SPAG8** | 83614 | 1.32E-02 | 2.22 |
| **SPARC** | 83644 | 2.33E-02 | 2.02 |
| **SPOCK3** | 84171 | 4.18E-02 | 2.25 |
| **SRD5A3** | 84521 | 5.29E-03 | 2.69 |
| **SRF** | 84555 | 6.09E-03 | 2.38 |
| **SSBP3** | 84840 | 2.68E-03 | 1.61 |
| **ST3GAL5** | 85034 | 2.89E-02 | 3.35 |
| **STK17A** | 85414 | 6.78E-03 | 1.99 |
| **SUV39H1** | 86144 | 2.93E-02 | 1.62 |
| **SYT9** | 86570 | 2.33E-02 | 2.54 |
| **TBCE** | 87449 | 4.29E-03 | 9.03 |
| **TESK2** | 88252 | 1.31E-02 | 1.45 |
| **TM9SF3** | 89574 | 2.93E-02 | 1.60 |
| **TNFAIP3** | 91321 | 1.50E-02 | 6.74 |
| **TRIM38** | 92737 | 4.77E-02 | 1.37 |
| **TSPAN18** | 93524 | 4.10E-02 | 1.73 |
| **TSPAN9** | 93586 | 3.10E-02 | 7.68 |
| **TTC16** | 93747 | 2.17E-02 | 1.83 |
| **TTC39C** | 93893 | 4.27E-02 | 1.58 |
| **TUBB4A** | 94196 | 3.01E-02 | 1.84 |
| **TWIST2** | 94369 | 4.17E-02 | 1.51 |
| **TXNIP** | 94468 | 3.97E-02 | 1.93 |
| **UBA1** | 94611 | 3.22E-02 | 1.60 |
| **UBL7** | 94960 | 3.96E-02 | 1.40 |
| **UBTD1** | 95041 | 1.45E-02 | 1.75 |
| **WASF2** | 96993 | 3.36E-02 | 2.48 |
| **WDR93** | 97447 | 3.84E-02 | 1.76 |
| **XKR6** | 97977 | 3.60E-02 | 2.29 |
| **ZBTB37** | 98557 | 3.70E-02 | 1.34 |
| **ZC3H4** | 98765 | 4.55E-02 | 5.03 |
| **ZFHX3** | 99127 | 8.00E-03 | 1.89 |
| **ZFP90** | 99244 | 4.29E-03 | 6.22 |
| **ZHX2** | 99370 | 4.38E-02 | 6.58 |
| **ZNF329** | 100271 | 4.10E-02 | 1.65 |
| **ZNF474** | 100742 | 1.90E-02 | 1.78 |
| **ZNF578** | 101169 | 7.12E-03 | 1.60 |
| **ZNF83** | 102147 | 3.14E-02 | 1.41 |
| **non-targeting** | 103174 | 7.12E-03 | 21.50 |
| **non-targeting** | 103334 | 3.01E-02 | 1.56 |
| **non-targeting** | 103418 | 3.13E-02 | 2.51 |
| **non-targeting** | 103596 | 3.36E-02 | 1.69 |
| **non-targeting** | 103650 | 2.74E-02 | 1.67 |
| **non-targeting** | 103960 | 4.53E-02 | 8.48 |
| **non-targeting** | 104192 | 4.03E-02 | 1.88 |

**Table S5.**
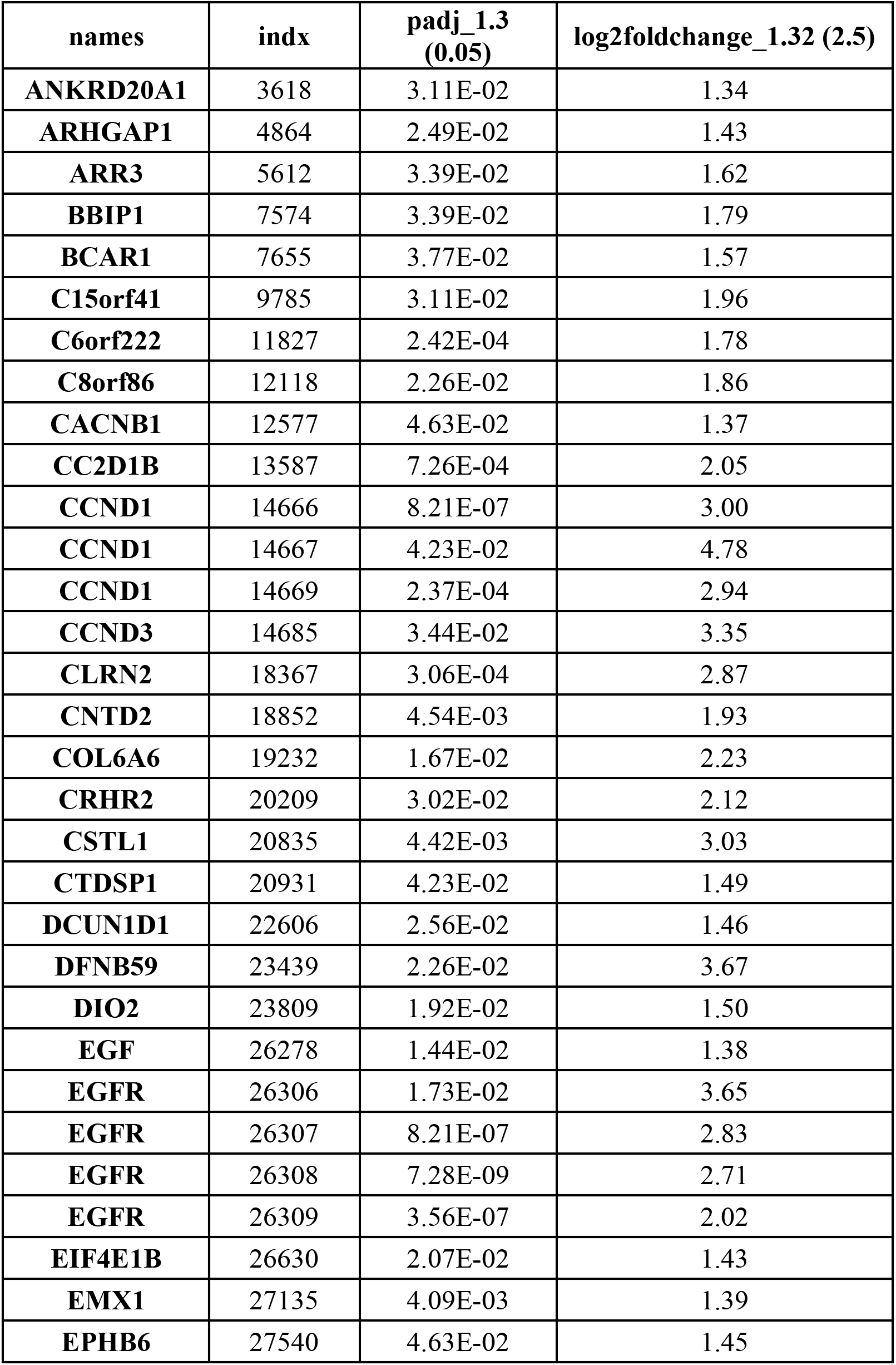
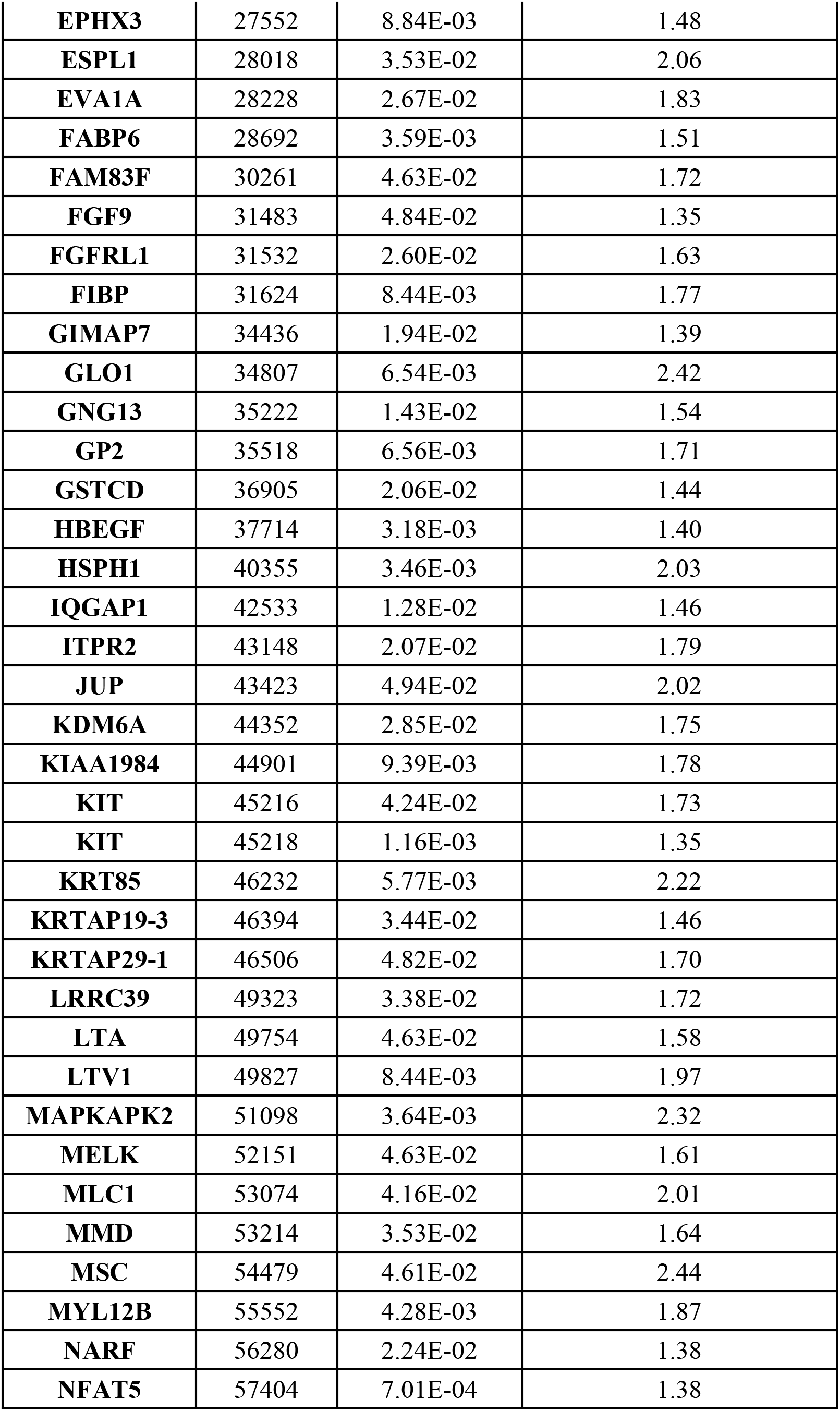
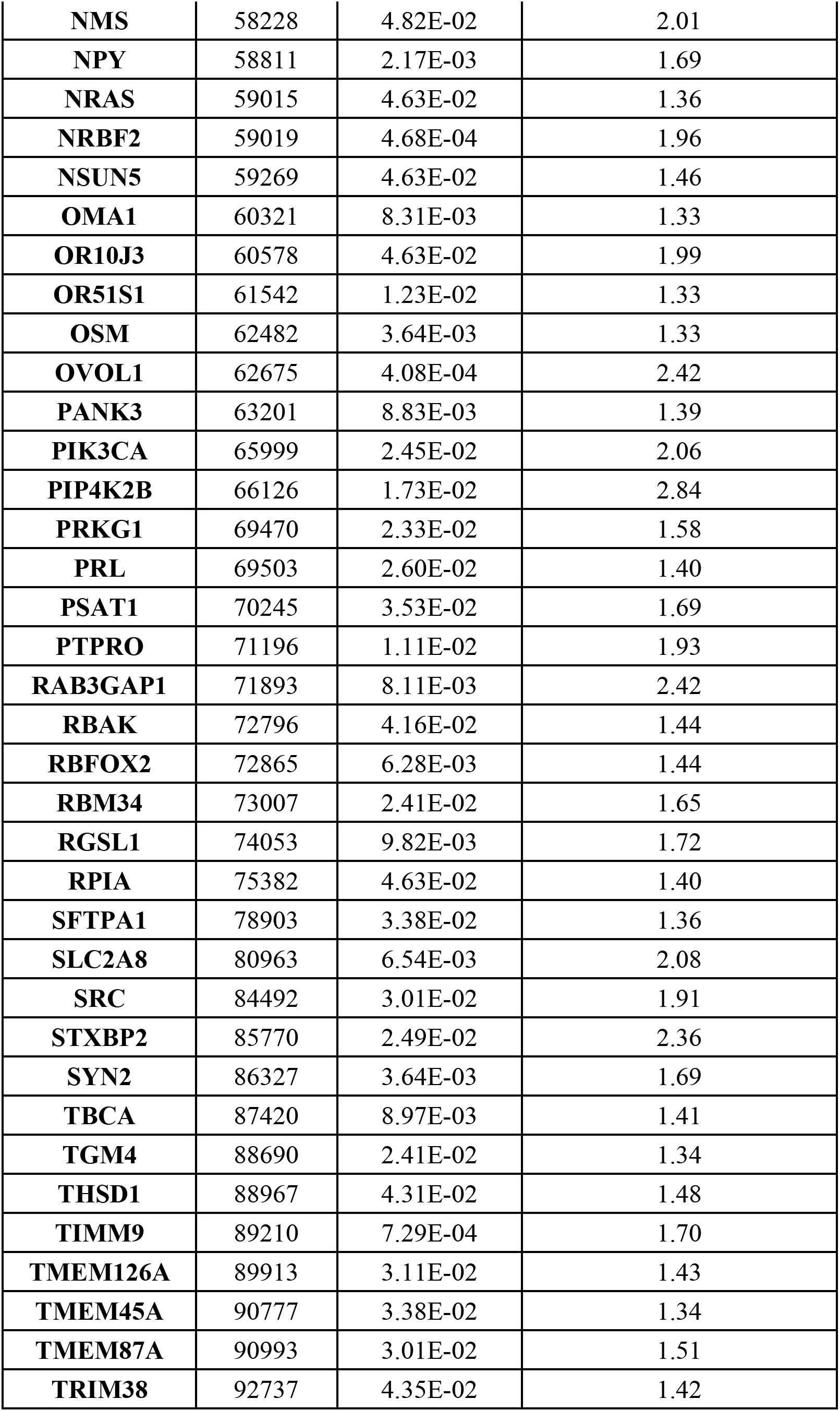
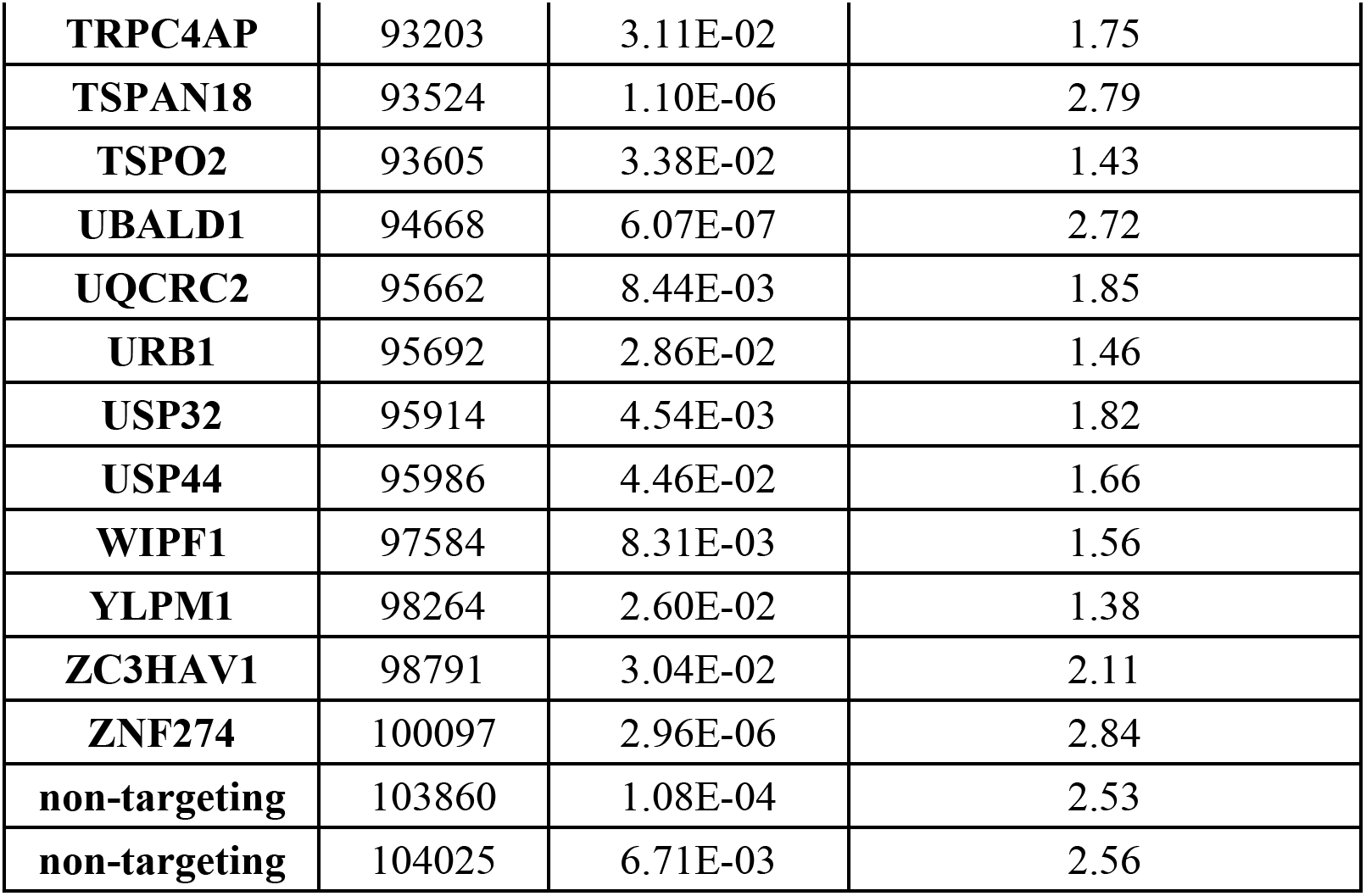
Significantly enriched gene comparison list for treatment comparison -Dox/+Tam vs. +Dox/+Tam.

**Table S6.** DisGeNet Enrichr disease gene set for -Dox/-Tam vs. -Dox/+Tam.

| **Term** | **Overlap** | **P-value** | **Adjusted P-value** | **Odds Ratio** | **Combined Score** | **Genes** |
| --- | --- | --- | --- | --- | --- | --- |
| **Brain Edema** | 3/63 | 0.0057 | 0.5426 | 8.8504 | 45.6962 | DBT;<br>CACNA1A;<br>GFAP |
| **Autism Spectrum Disorders** | 9/572 | 0.0058 | 0.5426 | 2.9139 | 15.0153 | NLGN4X; GRP;<br>CACNA1A;<br>ZSWIM6;<br>DGCR6; SOD3;<br>GIGYF2;<br>SSTR4; GFAP |
| **Clumsiness - motor delay** | 2/21 | 0.0064 | 0.5426 | 18.5058 | 93.4567 | ZSWIM6;<br>SSTR4 |
| **Dystonia Musculorum Deformans** | 2/22 | 0.0070 | 0.5426 | 17.5796 | 87.1697 | CIZ1;<br>CACNA1A |
| **Dystonia Disorders** | 4/135 | 0.0077 | 0.5426 | 5.4340 | 26.4400 | CIZ1; DBT;<br>CACNA1A;<br>TOR1AIP2 |
| **Dermatologic disorders** | 7/406 | 0.0091 | 0.5426 | 3.1654 | 14.8843 | TNFRSF17;<br>SUCLG1;<br>GATA3; SOD3;<br>GEM; BLVRA;<br>GFAP |
| **Machado-Joseph Disease** | 3/76 | 0.0096 | 0.5426 | 7.2696 | 33.7731 | CACNA1A;<br>BBC3; GFAP |
| **Aortic valve disorder** | 2/28 | 0.0112 | 0.5426 | 13.5187 | 60.6788 | BGN; SMAD6 |
| **Phenylketonurias** | 2/29 | 0.0120 | 0.5426 | 13.0174 | 57.5468 | CACNA1A;<br>GFAP |
| **Aortic valve calcification** | 2/32 | 0.0145 | 0.5426 | 11.7139 | 49.5699 | NPR1;SMAD6 |

**Table S7.** DisGeNet Enrichr disease gene set for -Dox/-Tam vs. +Dox/-Tam.

| Term | Overlap | P-value | Adjusted P-value | Odds Ratio | Combined Score | Genes |
| --- | --- | --- | --- | --- | --- | --- |
| Sinonasal undifferentiated carcinoma | 3/11 | 8.2E-06 | 1.4E-02 | 1.0E+02 | 1.2E+03 | NUTM1; ASCL1; EGFR |
| Supratentorial Embryonal Tumor, Not Otherwise Specified | 3/14 | 1.8E-05 | 1.5E-02 | 7.5E+01 | 8.2E+02 | DLC1; ASCL1; EGFR |
| Neuroectodermal Tumor, Primitive | 5/114 | 6.9E-05 | 4.0E-02 | 1.3E+01 | 1.2E+02 | MDK; SERPINF1; KDR; ASCL1; EGFR |
| Uremia | 4/67 | 1.2E-04 | 5.0E-02 | 1.8E+01 | 1.6E+02 | IL15; KDR; AGER; EGFR |
| Hormone refractory prostate cancer | 5/135 | 1.5E-04 | 5.3E-02 | 1.1E+01 | 9.5E+01 | PLAG1; KDR; ANXA7; AGER; EGFR |
| Secondary malignant neoplasm of lung | 10/683 | 2.3E-04 | 6.2E-02 | 4.4E+00 | 3.7E+01 | FERMT1; GPNMB; IL15; MDK; SERPINF1; DLC1; KDR; GRB2; AGER; EGFR |
| Medullary carcinoma of thyroid | 6/239 | 2.8E-04 | 6.2E-02 | 7.3E+00 | 6.0E+01 | CALCB; GRAP2; KDR; GRB2; ASCL1; EGFR |
| Subacute Bacterial Endocarditis | 2/7 | 2.9E-04 | 6.2E-02 | 1.1E+02 | 8.9E+02 | ASCL1; EGFR |
| Embryonal Rhabdomyosarcoma | 4/91 | 3.8E-04 | 6.6E-02 | 1.3E+01 | 1.0E+02 | GRAP2; CIB2; AGER; EGFR |
| Avascular retina | 2/8 | 3.8E-04 | 6.6E-02 | 9.1E+01 | 7.2E+02 | SERPINF1; KDR |

**Table S8.** DisGeNet Enrichr disease gene set for +Dox/-Tam vs. +Dox/+Tam.

| Term | Overlap | P-value | Adjusted P-value | Odds Ratio | Combined Score | Genes |
| --- | --- | --- | --- | --- | --- | --- |
| Germ Cell Cancer | 4/36 | 4.20E-04 | 0.3846 | 12.8051 | 99.5562 | ATF7IP;<br>KIT; EGFR;<br>FGF4 |
| stage, non-small cell lung cancer | 5/66 | 4.84E-04 | 0.3846 | 8.4281 | 64.3367 | ITGAM;<br>CCND1;<br>FAS;<br>CXCL5;<br>EGFR |
| Childhood Acute Myeloid Leukemia | 6/102 | 5.20E-04 | 0.3846 | 6.4486 | 48.7589 | KIT; FAS;<br>GATA2;<br>GATA1;<br>CBFA2T3;<br>RPRM |
| Primary Carcinoma | 4/43 | 8.34E-04 | 0.3846 | 10.5030 | 74.4543 | ITGAM;<br>CCND1;<br>KIT; EGFR |
| Intestinal Neoplasms | 7/158 | 9.86E-04 | 0.3846 | 4.7948 | 33.1893 | ITGAM;<br>CCND1;<br>KIT; PRR15;<br>FAS;<br>INSM1;<br>EGFR |
| keratoacanthoma | 4/48 | 0.00127 | 0.3846 | 9.3071 | 62.0954 | ZHX2;<br>CCND1;<br>FAS; DMKN |
| Male Germ Cell Tumor | 2/6 | 0.00141 | 0.3846 | 50.7667 | 333.2138 | KIT; FAS |
| Carcinosarcoma | 4/50 | 0.00148 | 0.3846 | 8.9016 | 58.0255 | CCND1;<br>KIT; EGFR;<br>MAGEA4 |
| Bronchioloalveolar Adenocarcinoma | 5/89 | 0.00187 | 0.3846 | 6.1133 | 38.3912 | ATF2;<br>DYRK2;<br>CCND1;<br>SFTPC;<br>EGFR |
| Non-small cell lung cancer stage III | 2/7 | 0.00196 | 0.3846 | 40.6113 | 253.1570 | CCND1;<br>EGFR |

**Table S9.**
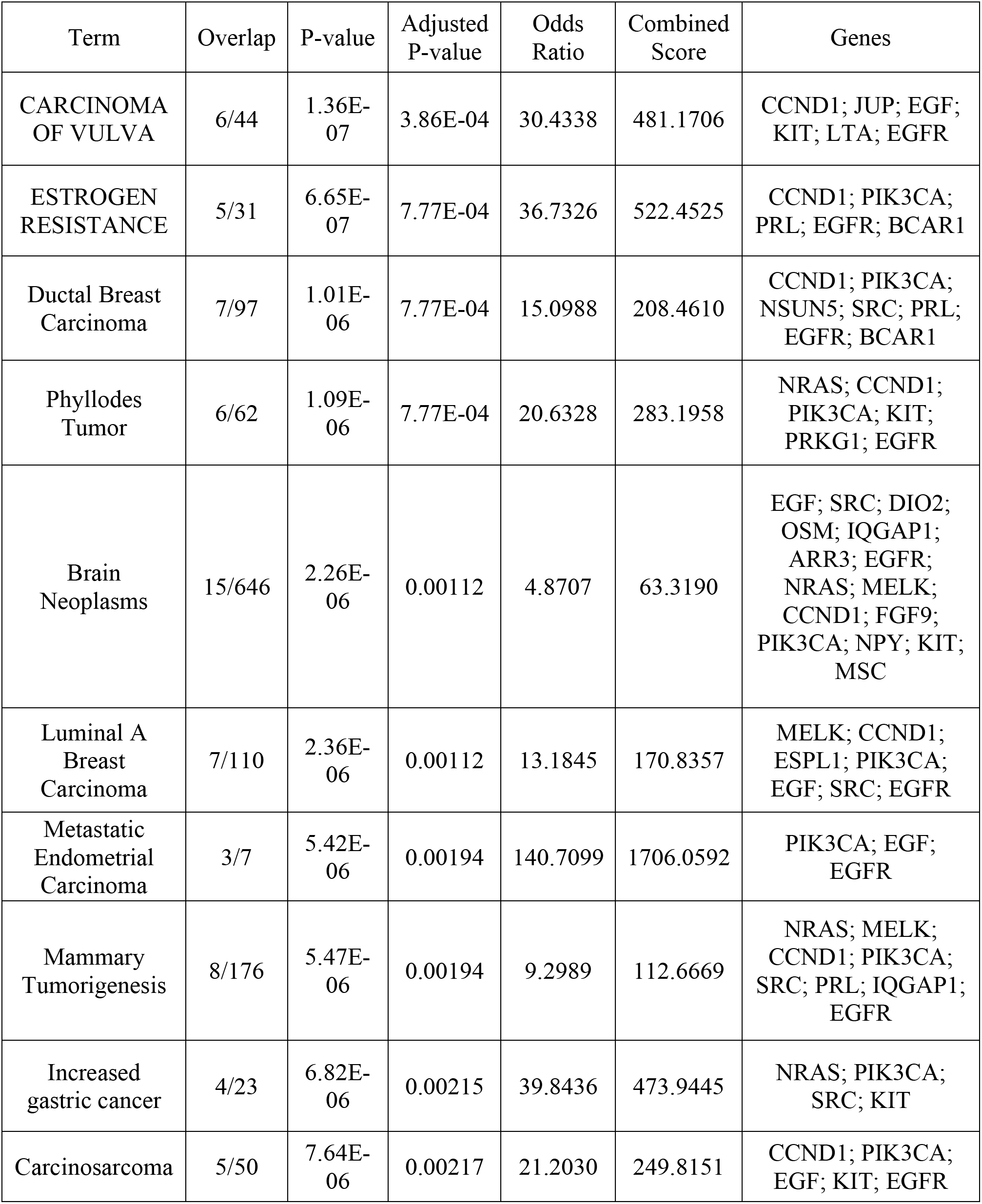
DisGeNet Enrichr disease gene set for -Dox/+Tam vs. +Dox/+Tam.

**Table S10.**
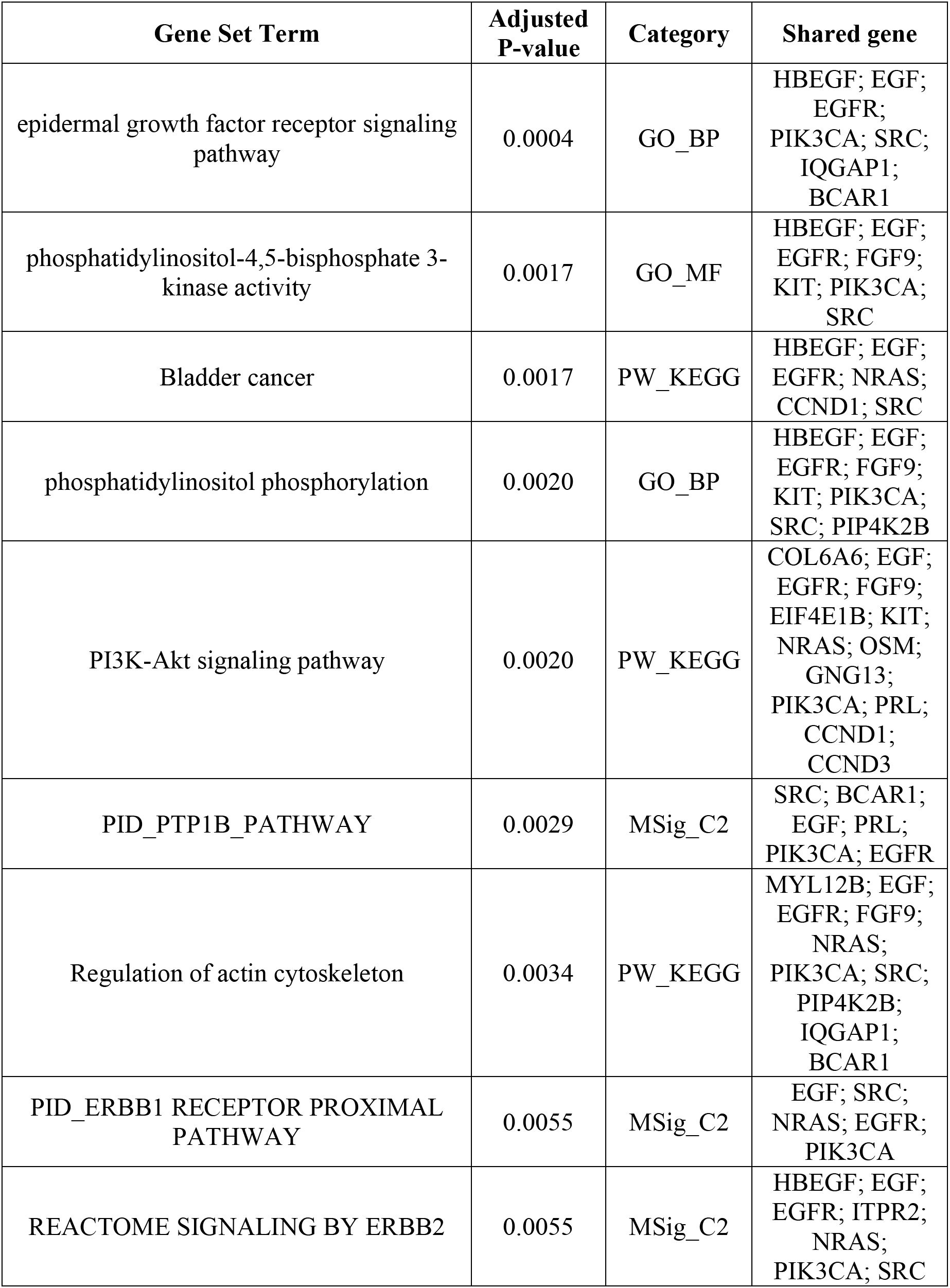
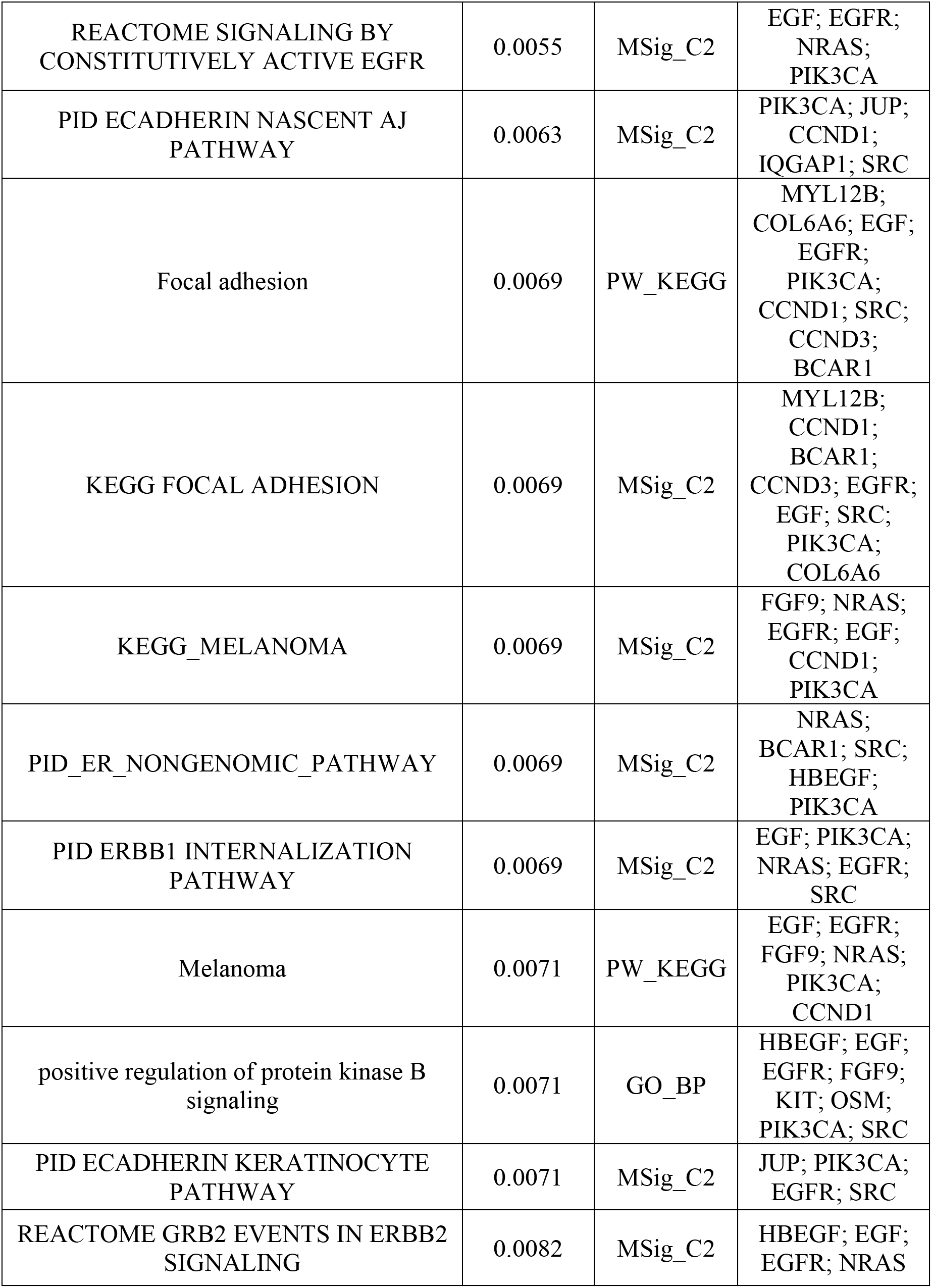
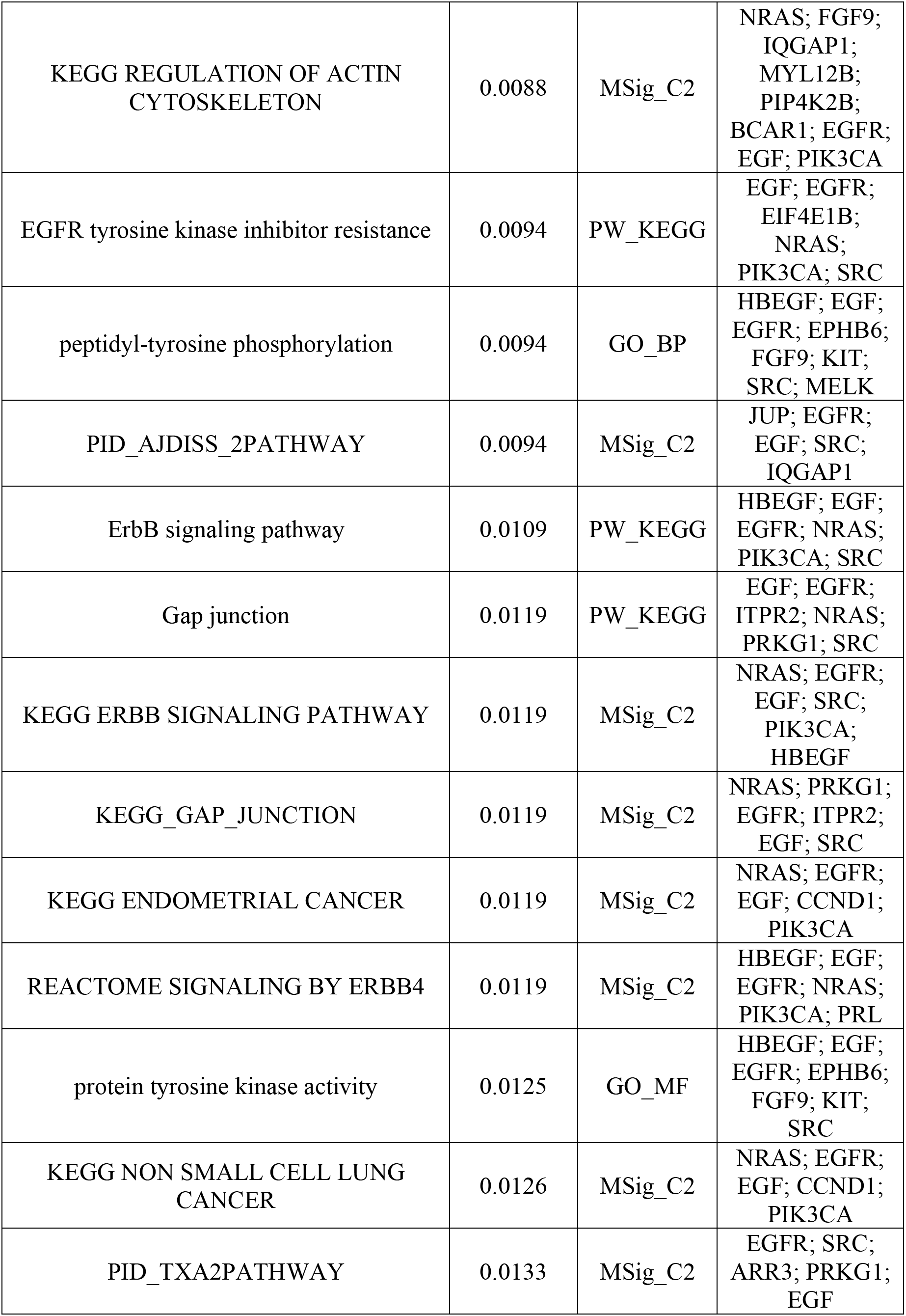
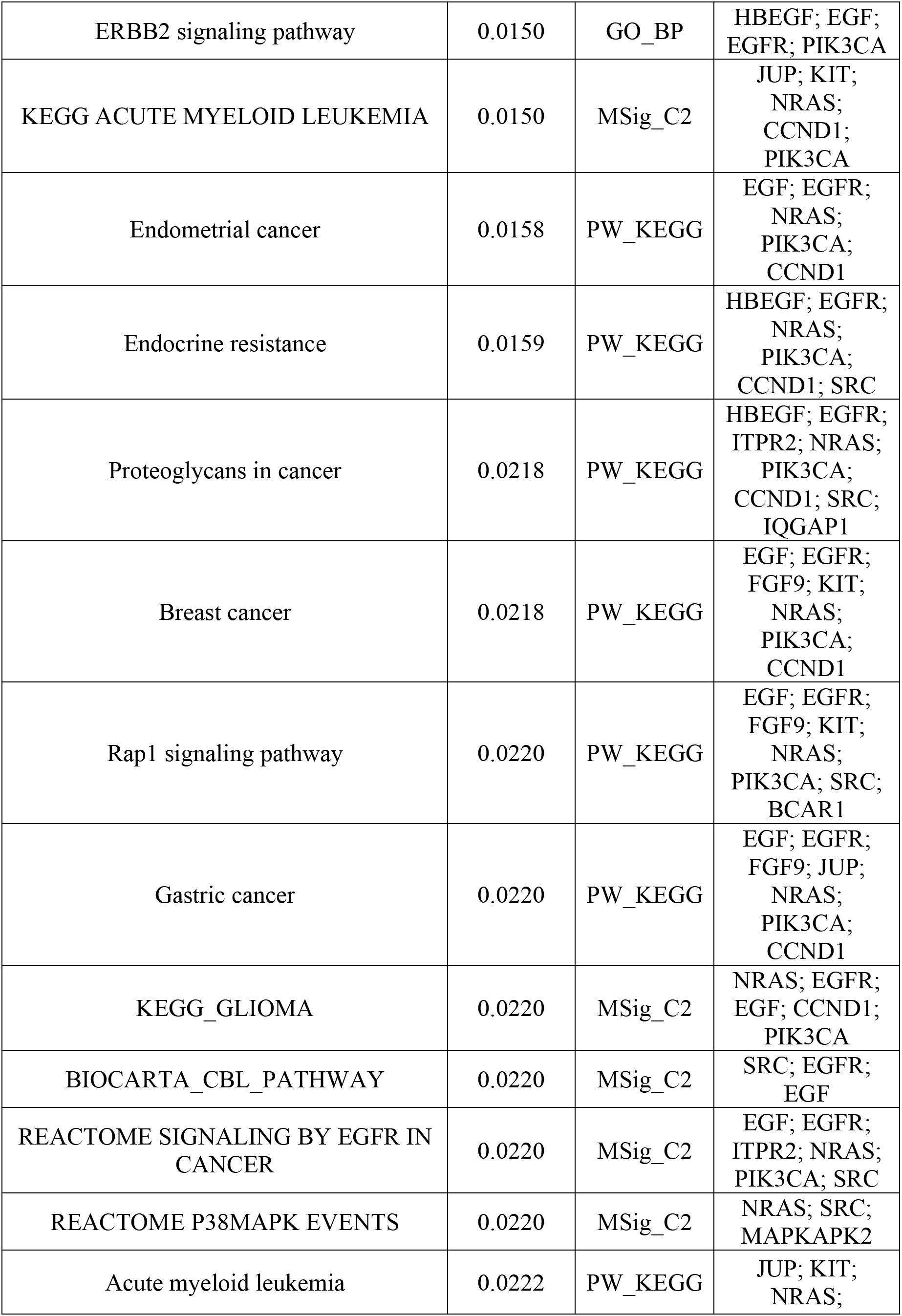
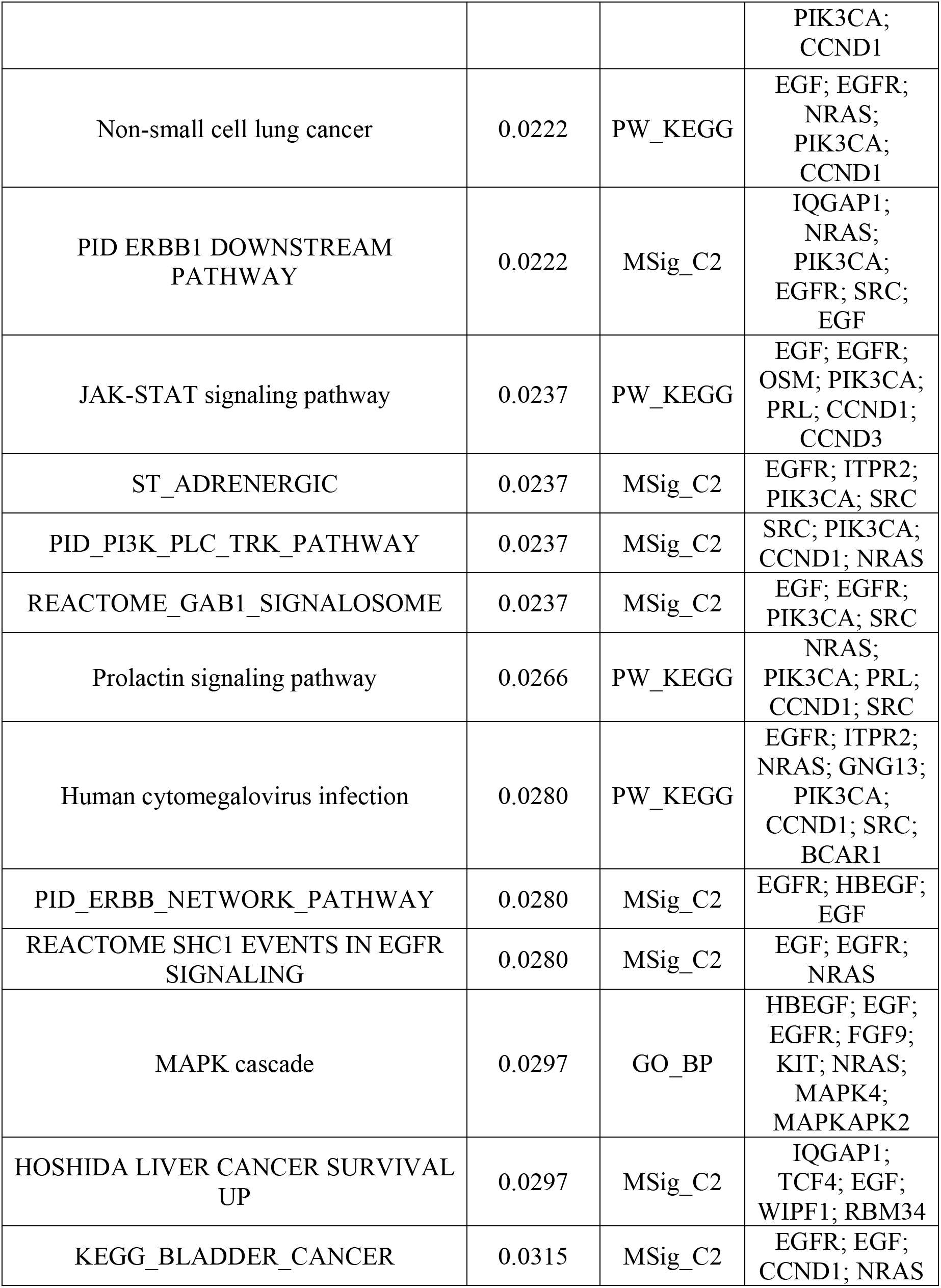
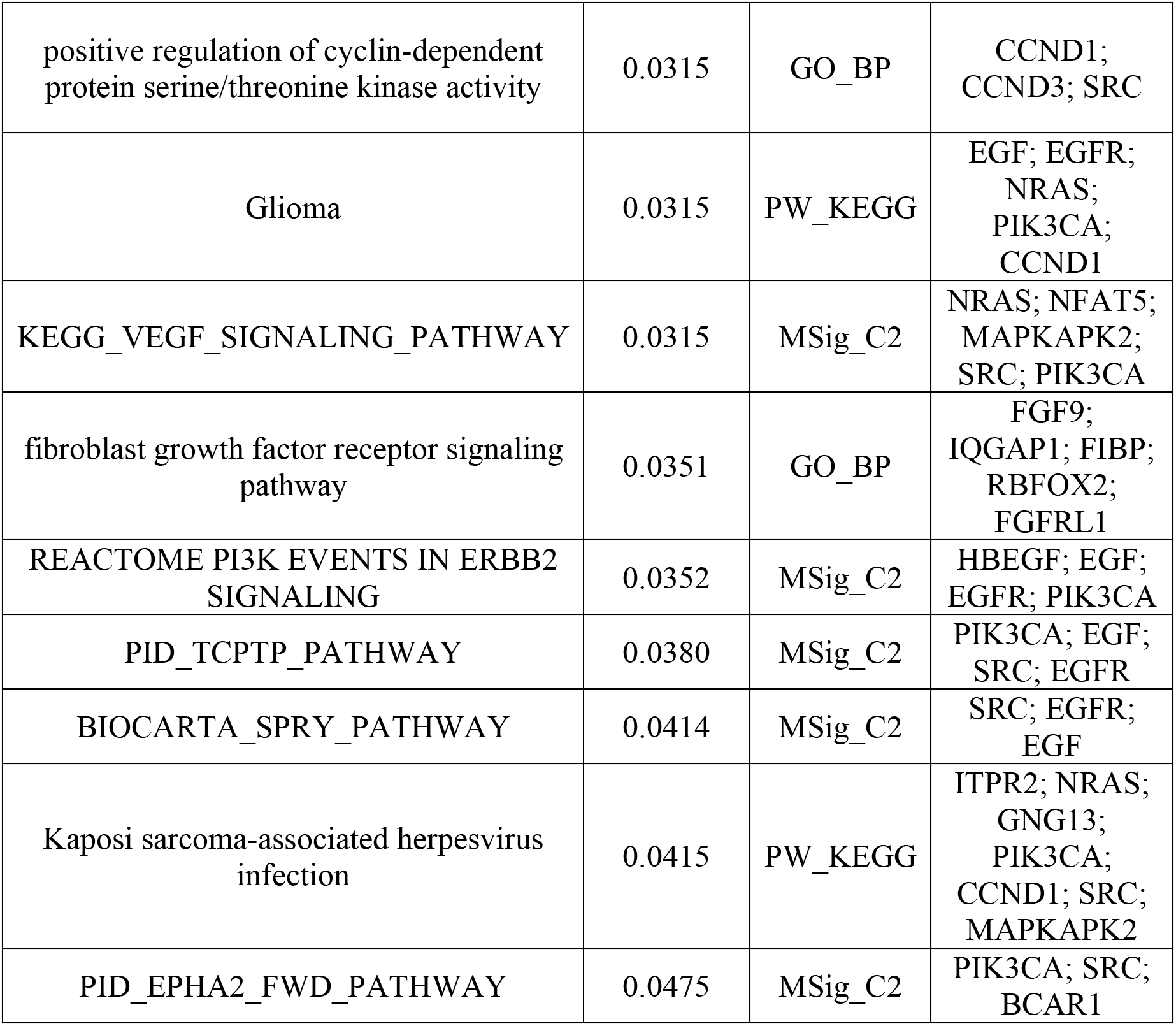
HTSanalyzeR2 enrichment map of gene set over-representation analysis (GSOA)

**Table S11.**
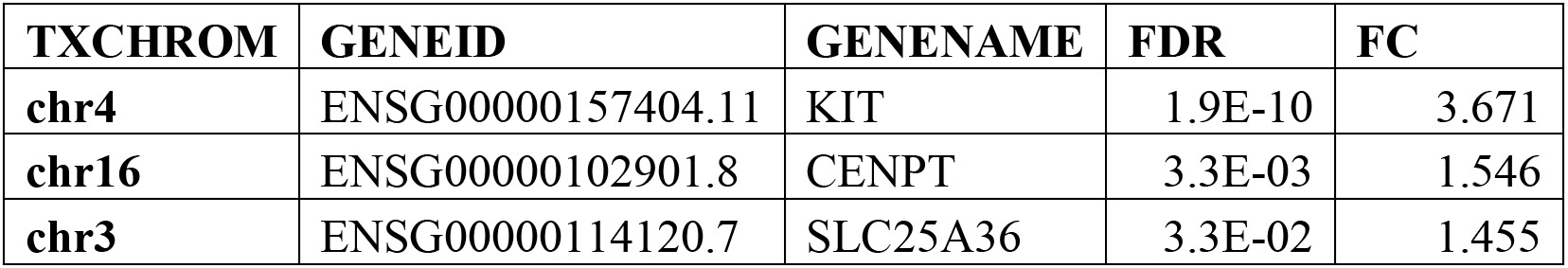
24hr Dox/Vehicle Treatment, PRO-seq.

| **TXCHROM** | **GENEID** | **GENENAME** | **FDR** | **FC** |
| --- | --- | --- | --- | --- |
| **chr4** | ENSG00000157404.11 | KIT | 1.9E-10 | 3.671 |
| **chr16** | ENSG00000102901.8 | CENPT | 3.3E-03 | 1.546 |
| **chr3** | ENSG00000114120.7 | SLC25A36 | 3.3E-02 | 1.455 |

**Table S12.**
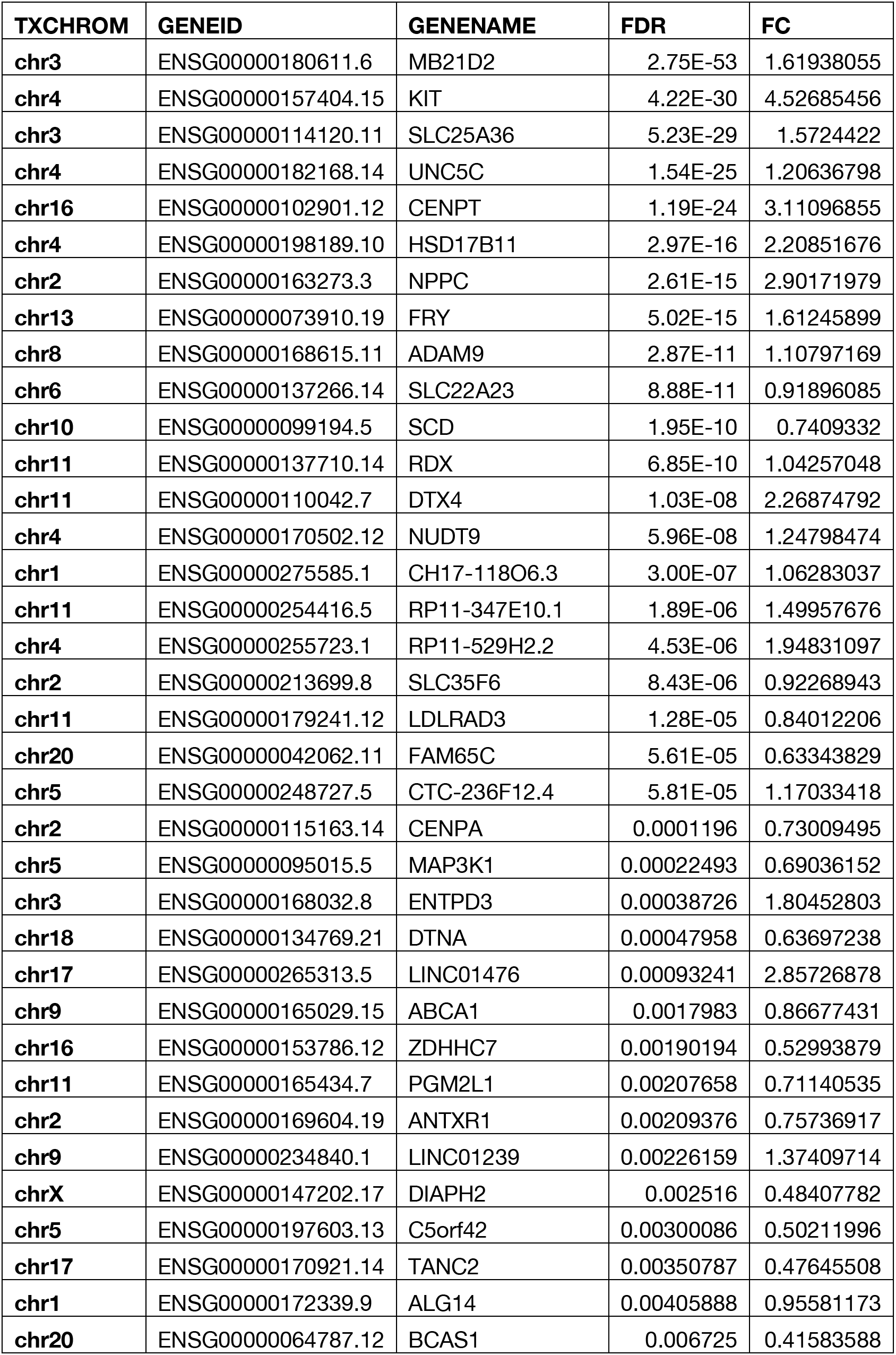
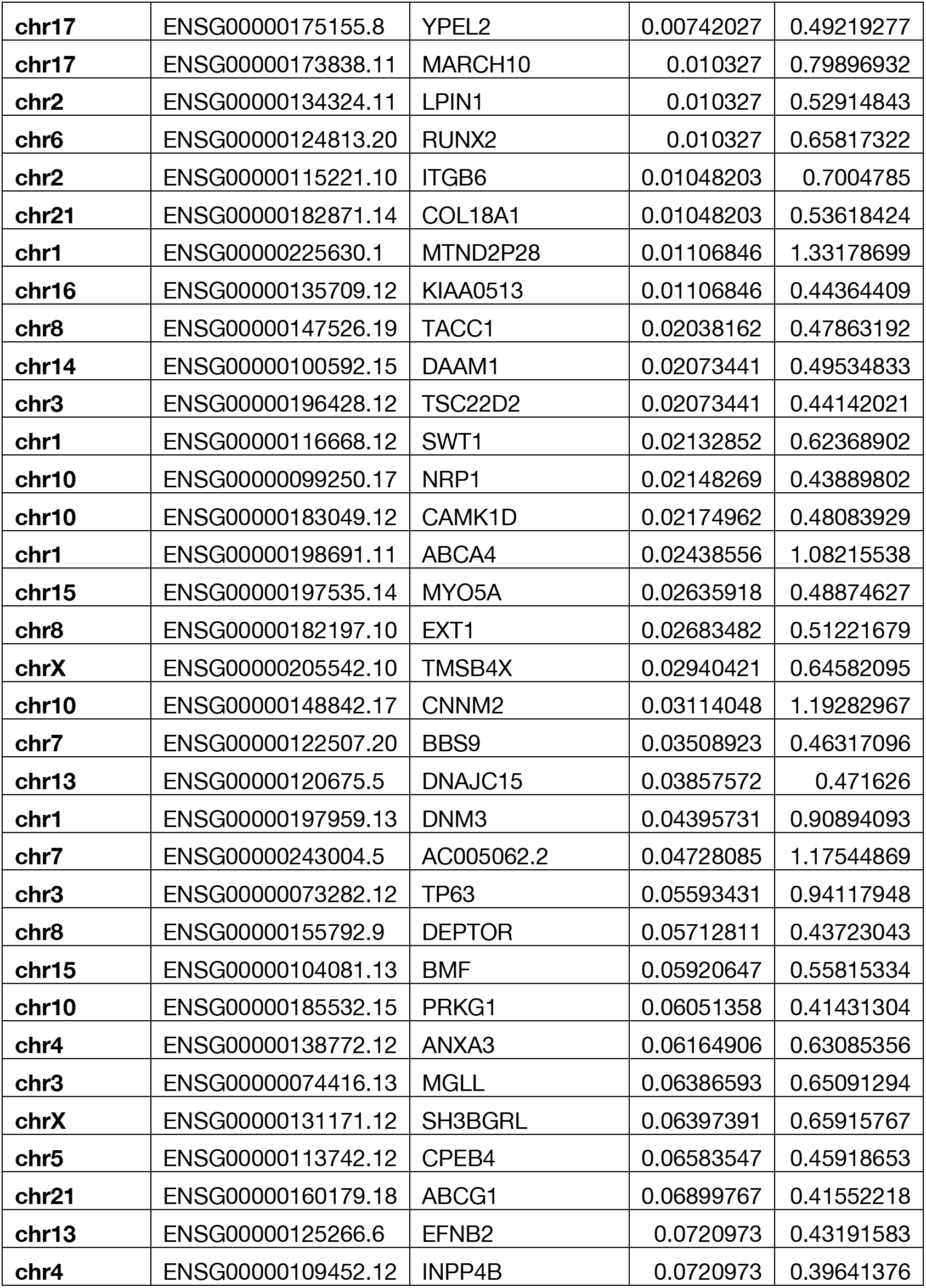
5-day Dox/Vehicle (+TAM) enriched gene set, p-value < 0.075, PROseq.

**Table S13.** KEGG 2021 Enriched Pathways using Enrichr.

| Term | Overlap | P-value | Adjusted P-value | Genes |
| --- | --- | --- | --- | --- |
| ABC transporters | 3/45 | 5.2E-04 | 4.1E-02 | ABCA1; ABCA4; ABCG1 |
| Regulation of actin cytoskeleton | 4/218 | 7.2E-03 | 2.8E-01 | DIAPH2; TMSB4X; RDX; ITGB6 |
| Regulation of lipolysis in adipocytes | 2/55 | 1.6E-02 | 3.8E-01 | PRKG1; MGLL |
| Glycerolipid metabolism | 2/61 | 1.9E-02 | 3.8E-01 | LPIN1; MGLL |
| Axon guidance | 3/182 | 2.6E-02 | 4.1E-01 | EFNB2; NRP1; UNC5C |
| Purine metabolism | 2/129 | 7.5E-02 | 4.8E-01 | NUDT9; ENTPD3 |
| Vascular smooth muscle contraction | 2/133 | 7.9E-02 | 4.8E-01 | NPPC; PRKG1 |
| Biosynthesis of unsaturated fatty acids | 1/27 | 9.0E-02 | 4.8E-01 | SCD |
| MicroRNAs in cancer | 3/310 | 9.5E-02 | 4.8E-01 | RDX; BMF; TP63 |
| Phospholipase D signaling pathway | 2/148 | 9.5E-02 | 4.8E-01 | DNM3; KIT |
| mTOR signaling pathway | 2/154 | 1.0E-01 | 4.8E-01 | DEPTOR; LPIN1 |
| cGMP-PKG signaling pathway | 2/167 | 1.2E-01 | 4.8E-01 | NPPC; PRKG1 |
| Tight junction | 2/169 | 1.2E-01 | 4.8E-01 | MAP3K1; RDX |
| Starch and sucrose metabolism | 1/36 | 1.2E-01 | 4.8E-01 | PGM2L1 |
| Various types of N-glycan biosynthesis | 1/39 | 1.3E-01 | 4.8E-01 | ALG14 |
| Fat digestion and absorption | 1/43 | 1.4E-01 | 4.8E-01 | ABCA1 |
| N-Glycan biosynthesis | 1/50 | 1.6E-01 | 4.8E-01 | ALG14 |
| Cholesterol metabolism | 1/50 | 1.6E-01 | 4.8E-01 | ABCA1 |
| Endocrine and other factor-regulated calcium reabsorption | 1/53 | 1.7E-01 | 4.8E-01 | DNM3 |
| Glycosaminoglycan biosynthesis | 1/53 | 1.7E-01 | 4.8E-01 | EXT1 |
| Lipid and atherosclerosis | 2/215 | 1.7E-01 | 4.8E-01 | ABCA1; ABCG1 |
| Pyrimidine metabolism | 1/56 | 1.8E-01 | 4.8E-01 | ENTPD3 |
| Human T-cell leukemia virus 1 infection | 2/219 | 1.8E-01 | 4.8E-01 | NRP1; MAP3K1 |
| Notch signaling pathway | 1/59 | 1.9E-01 | 4.8E-01 | DTX4 |
| Long-term depression | 1/60 | 1.9E-01 | 4.8E-01 | PRKG1 |
| Thermogenesis | 2/232 | 1.9E-01 | 4.8E-01 | MGLL; PRKG1 |
| Acute myeloid leukemia | 1/67 | 2.1E-01 | 4.8E-01 | KIT |
| Central carbon metabolism in cancer | 1/70 | 2.2E-01 | 4.8E-01 | KIT |
| RIG-I-like receptor signaling pathway | 1/70 | 2.2E-01 | 4.8E-01 | MAP3K1 |
| Inositol phosphate metabolism | 1/73 | 2.3E-01 | 4.8E-01 | INPP4B |
| PPAR signaling pathway | 1/74 | 2.3E-01 | 4.8E-01 | SCD |
| Glioma | 1/75 | 2.3E-01 | 4.8E-01 | CAMK1D |
| Arrhythmogenic right ventricular cardiomyopathy | 1/77 | 2.4E-01 | 4.8E-01 | ITGB6 |
| Bacterial invasion of epithelial cells | 1/77 | 2.4E-01 | 4.8E-01 | DNM3 |
| Synaptic vesicle cycle | 1/78 | 2.4E-01 | 4.8E-01 | DNM3 |
| ECM-receptor interaction | 1/88 | 2.7E-01 | 4.8E-01 | ITGB6 |
| Gap junction | 1/88 | 2.7E-01 | 4.8E-01 | PRKG1 |
| Hypertrophic cardiomyopathy | 1/90 | 2.7E-01 | 4.8E-01 | ITGB6 |
| MAPK signaling pathway | 2/294 | 2.7E-01 | 4.8E-01 | MAP3K1; KIT |
| Salivary secretion | 1/93 | 2.8E-01 | 4.8E-01 | PRKG1 |
| GnRH signaling pathway | 1/93 | 2.8E-01 | 4.8E-01 | MAP3K1 |
| Dilated cardiomyopathy | 1/96 | 2.9E-01 | 4.8E-01 | ITGB6 |
| Circadian entrainment | 1/97 | 2.9E-01 | 4.8E-01 | PRKG1 |
| Phosphatidylinositol signaling system | 1/97 | 2.9E-01 | 4.8E-01 | INPP4B |
| Aldosterone synthesis and secretion | 1/98 | 2.9E-01 | 4.8E-01 | CAMK1D |
| Glycerophospholipid metabolism | 1/98 | 2.9E-01 | 4.8E-01 | LPIN1 |
| Hematopoietic cell lineage | 1/99 | 2.9E-01 | 4.8E-01 | KIT |
| Progesterone-mediated oocyte maturation | 1/100 | 3.0E-01 | 4.8E-01 | CPEB4 |
| Melanogenesis | 1/101 | 3.0E-01 | 4.8E-01 | KIT |
| Protein digestion and absorption | 1/103 | 3.0E-01 | 4.8E-01 | COL18A1 |
| Parathyroid hormone synthesis, secretion and action | 1/106 | 3.1E-01 | 4.8E-01 | RUNX2 |
| Neurotrophin signaling pathway | 1/119 | 3.4E-01 | 5.0E-01 | MAP3K1 |
| Growth hormone synthesis, secretion and action | 1/119 | 3.4E-01 | 5.0E-01 | MAP3K1 |
| AMPK signaling pathway | 1/120 | 3.4E-01 | 5.0E-01 | SCD |
| PI3K-Akt signaling pathway | 2/354 | 3.5E-01 | 5.0E-01 | KIT;ITGB6 |
| Platelet activation | 1/124 | 3.5E-01 | 5.0E-01 | PRKG1 |
| Oocyte meiosis | 1/129 | 3.6E-01 | 5.1E-01 | CPEB4 |
| Autophagy | 1/137 | 3.8E-01 | 5.1E-01 | DEPTOR |
| Fluid shear stress and atherosclerosis | 1/139 | 3.9E-01 | 5.1E-01 | NPPC |
| Ubiquitin mediated proteolysis | 1/140 | 3.9E-01 | 5.1E-01 | MAP3K1 |
| Breast cancer | 1/147 | 4.0E-01 | 5.2E-01 | KIT |
| Retrograde endocannabinoid signaling | 1/148 | 4.1E-01 | 5.2E-01 | MGLL |
| Oxytocin signaling pathway | 1/154 | 4.2E-01 | 5.2E-01 | CAMK1D |
| Hepatitis B | 1/162 | 4.3E-01 | 5.4E-01 | MAP3K1 |
| Wnt signaling pathway | 1/166 | 4.4E-01 | 5.4E-01 | DAAM1 |
| NOD-like receptor signaling pathway | 1/181 | 4.7E-01 | 5.6E-01 | ANTXR1 |
| Transcriptional misregulation in cancer | 1/192 | 4.9E-01 | 5.7E-01 | RUNX2 |
| Pathogenic Escherichia coli infection | 1/197 | 5.0E-01 | 5.7E-01 | MYO5A |
| Focal adhesion | 1/201 | 5.1E-01 | 5.7E-01 | ITGB6 |
| Epstein-Barr virus infection | 1/202 | 5.1E-01 | 5.7E-01 | ENTPD3 |
| Proteoglycans in cancer | 1/205 | 5.1E-01 | 5.7E-01 | RDX |
| Rap1 signaling pathway | 1/210 | 5.2E-01 | 5.7E-01 | KIT |
| Coronavirus disease | 1/232 | 5.6E-01 | 6.0E-01 | NRP1 |
| Ras signaling pathway | 1/232 | 5.6E-01 | 6.0E-01 | KIT |
| Calcium signaling pathway | 1/240 | 5.7E-01 | 6.0E-01 | CAMK1D |
| Endocytosis | 1/252 | 5.9E-01 | 6.1E-01 | DNM3 |
| Human papillomavirus infection | 1/331 | 6.9E-01 | 7.1E-01 | ITGB6 |
| Olfactory transduction | 1/440 | 7.9E-01 | 8.0E-01 | PRKG1 |
| Pathways in cancer | 1/531 | 8.5E-01 | 8.5E-01 | KIT |

**Table S14.**
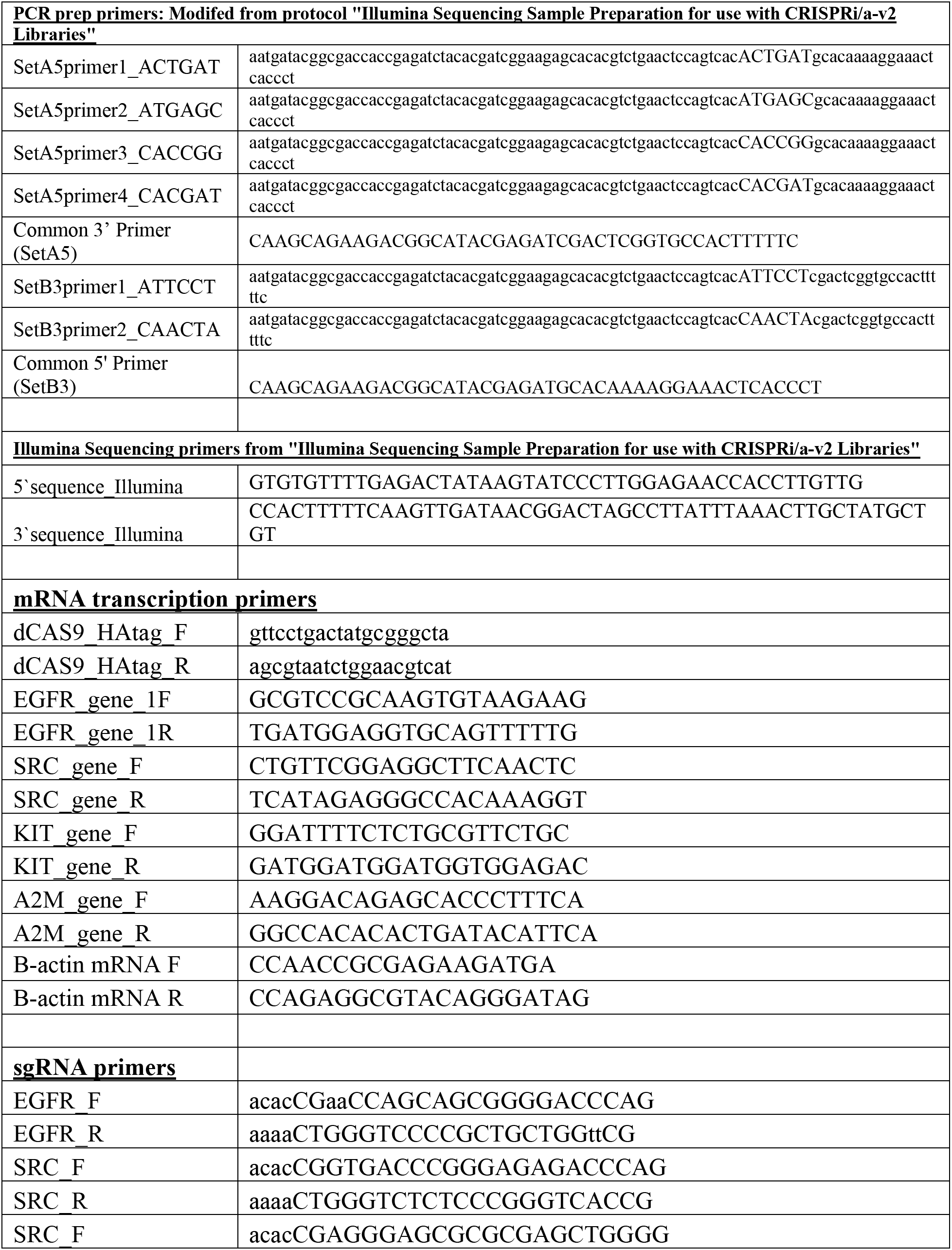
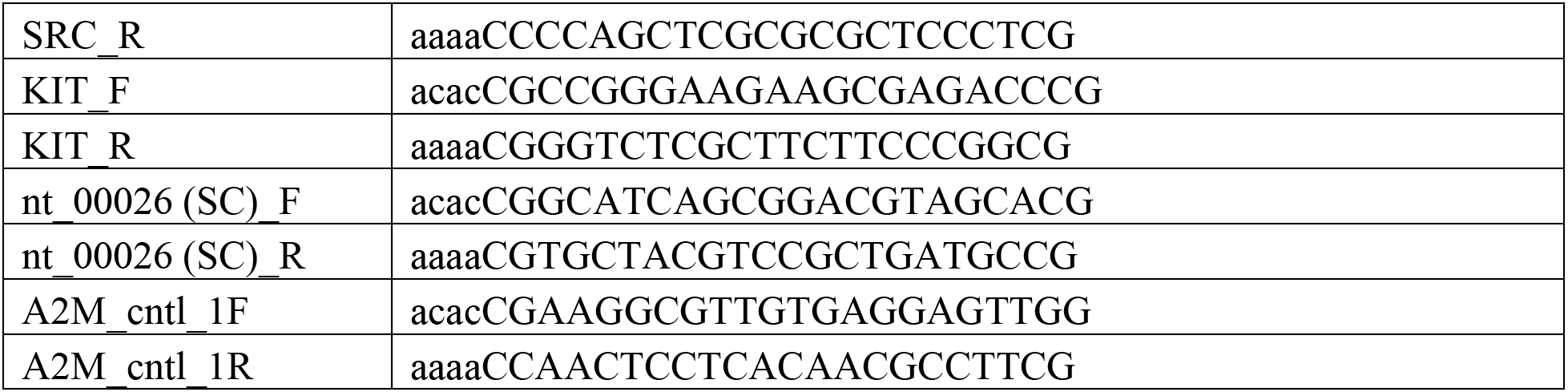
Primers.

## Notes

### Competing Interest Statement

The authors have declared no competing interest.

